# CyAbrB2 is a nucleoid-associated protein in *Synechocystis* controlling hydrogenase expression during fermentation

**DOI:** 10.1101/2023.04.28.538649

**Authors:** Ryo Kariyazono, Takashi Osanai

## Abstract

The *hox* operon in *Synechocystis* sp. PCC 6803, encoding bidirectional hydrogenase responsible for H_2_ production, is transcriptionally upregulated under microoxic conditions. Although several regulators for *hox* transcription have been identified, their dynamics and higher-order DNA structure of *hox* region in microoxic conditions remain elusive. We focused on key regulators for the *hox* operon: cyAbrB2, a conserved regulator in cyanobacteria, and SigE, an alternative sigma factor. Chromatin immunoprecipitation-sequencing revealed that cyAbrB2 binds to the *hox* promoter region under aerobic conditions, with its binding being flattened in microoxic conditions. Concurrently, SigE exhibited increased localization to the *hox* promoter under microoxic conditions. Genome-wide analysis revealed that cyAbrB2 binds broadly to AT-rich genome regions and represses gene expression. Moreover, we demonstrated the physical interactions of the *hox* promoter region with its distal genomic loci. Both the transition to microoxic conditions and the absence of cyAbrB2 influenced the chromosomal interaction. From these results, we propose that cyAbrB2 is a cyanobacterial nucleoid-associated protein (NAP), modulating chromosomal conformation, which blocks RNA polymerase from the *hox* promoter in aerobic conditions. We further infer that cyAbrB2, with altered localization pattern upon microoxic conditions, modifies chromosomal conformation in microoxic conditions, which allows SigE-containing RNA polymerase to access the *hox* promoter. The coordinated actions of this NAP and the alternative sigma factor are crucial for the proper *hox* expression in microoxic conditions. Our results highlight the impact of cyanobacterial chromosome conformation and NAPs on transcription, which have been insufficiently investigated.

## Introduction

Cyanobacteria perform fermentation, using glycolytic products as electron acceptors. (1). Cyanobacteria have multiple fermentation pathways according to the environment. For example, the freshwater living cyanobacterium *Synechocystis* sp. PCC 6803 (hereafter referred to as *Synechocystis*) produces acetate, lactate, dicarboxylic acids, and hydrogen (1, 2).

Hydrogen is generated in quantities comparable to lactate and dicarboxylic acids as the result of electron acceptance in the dark microoxic condition (3, 4). Bidirectional hydrogenase is a key enzyme for H_2_ production from protons (5) and is commonly found in cyanobacteria (6). Cyanobacterial hydrogenase comprises five subunits (HoxEFUHY) containing nickel and Fe-S clusters (7). This enzyme can utilize NADH, reduced ferredoxin, and flavodoxin as substrates (8). Hydrogenase mainly receives reduced ferredoxin from pyruvate-ferredoxin oxidoreductase (PFOR) in the microoxic condition (8, 9). Although hydrogenase and PFOR are O_2_ sensitive, they can work under aerobic conditions (10–12). Therefore, uncontrolled expression of *hox* operon and *nifJ* (coding gene of PFOR) may hamper metabolism under photosynthetic conditions. Furthermore, genetic manipulations on *Synechocystis* have demonstrated that modulating the expression of certain enzymes including hydrogenase can alter fermentative metabolic flow (3, 4, 13). This provides evidence that transcription regulates the fermentative pathway. Thus, transcriptional regulation in response to the environment is essential for optimal energy cost performance.

Promoter recognition by RNA polymerases is an essential step in transcriptional regulation. Sigma factors, subunits of RNA polymerase, recognize core promoter sequences. Transcription factors can also bind to promoter regions to suppress or promote RNA polymerase transcription. As well as recruitment or blocking of RNA polymerase, some transcriptional factors, known as Nucleoid-associated proteins (NAPs), modulate chromosomal conformation to regulate transcription (14). NAPs are common in bacteria, but cyanobacterial NAPs remain unidentified, and higher-order DNA structure in cyanobacteria is yet to be shown. A recent study suggested that the manipulation of chromosomal supercoiling impacts transcriptional properties in cyanobacteria (15). There is room for consideration of NAPs modulating chromosomal conformation and regulating expression in cyanobacteria.

In *Synechosysits*, the coding genes of HoxEFUHY form a single operon (sll1220–1226), while PFOR is encoded in the *nifJ* (sll0741) gene. Both *hox* and *nifJ* operons are highly expressed under microoxic conditions (16). Genetic analysis has revealed that multiple global transcriptional regulators control *hox* and *nifJ* expression. Sigma factor SigE (Sll1689) promotes the expression of *hox* and *nifJ* operons (17, 18), while transcription factor cyAbrB2 (Sll0822) represses them (19, 20). Positive regulators for the *hox* operon include LexA (Sll1626) and cyAbrB1 (Sll0359) (21–23).

SigE, an alternative sigma factor, controls the expression of genes involved in glycogen catabolism and glycolysis, as well as PFOR/*nifJ* and hydrogenase (17). SigE shows a high amino acid sequence similarity with the primary sigma factor SigA, which is responsible for transcribing housekeeping and photosynthetic genes (24). A ChIP-seq study revealed that, while most SigE-binding sites are the same as SigA, SigE exclusively occupies the promoters of glycogen catabolism and glycolysis (25).

CyAbrB2 and its homolog cyAbrB1 are transcription factors highly conserved in cyanobacteria. For example, cyAbrB homologs in *Anabaena* sp. PCC7120 is involved in heterocyst formation (26). CyAbrB2 in *Synechocystis* regulates the expression of several genes involved in carbon metabolism, nitrogen metabolism, and cell division (20, 27, 28). CyAbrB2 binds to the *hox* promoter *in vitro* and represses its expression *in vivo* (19). CyAbrB1, an essential gene, physically interacts with the cyAbrB2 protein (29) and binds the *hox* promotor region *in vitro* to promote its expression (21).

To explore the dynamics of those transcription factors governing the expression of *hox* operon, we conducted a time-course analysis of the transcriptome and ChIP-seq of SigE and cyAbrB2. Our ChIP-seq and transcriptome analysis showed the NAPs-like nature of cyAbrB2, which prompted us to conduct a chromosomal conformation capture assay. 3C analysis explored the physical interaction between the *hox* promoter region and its downstream and upstream genomic region in the aerobic condition, and some loci changed interaction frequency upon entry to the microoxic condition. Furthermore, some interactions in the Δ*cyabrB2* mutant were different from those of the wildtype. From those experiments, we propose that cyAbrB2 modulates chromosomal conformation like NAPs, allowing access to the SigE-containing RNA polymerase complex on the *hox* promoter, by which the *hox* operon achieves distinct expression dynamics. Chromosomal conformation of bacteria is a growing area of interest, and the findings of this study have brought insight into the transcriptional regulation of cyanobacteria.

## Results

### Transcriptomes on entry to dark microoxic conditions

To understand transcriptional regulation under microoxic conditions, we conducted a time-course transcriptome capturing light aerobic and dark microoxic conditions at 1-, 2-, and 4-hours time points (Figure 1A). Gene set enrichment analysis based on KEGG pathway revealed that many biological pathways, including photosynthesis and respiration (oxidative phosphorylation), were downregulated by the transition to dark microoxic conditions from light aerobic conditions (Figure 1B). Upregulated pathways included butanoate metabolism and two-component systems. The enrichment in the butanoate metabolism pathway indicates the upregulation of genes involved in carbohydrate metabolism. We further classified genes according to their expression dynamics. Within 1 hour of switching from aerobic to microoxic conditions, the expression levels of 508 genes increased more than 2-fold. Furthermore, genes with increased expression levels were classified into four groups based on the time course (Figures 1C and S1). Of the 508 genes, 28 were termed “transiently upregulated genes” due to their decreased expression upon the comparison of 1- and 4-hours incubation under microoxic conditions (log2 fold change < −0.5), and 119 were termed “continuously upregulated genes.”, which continuously increased between 1- and 4-hours incubation under microoxic conditions (log2-fold change > 0.5). Other than 508 genes 2-fold upregulated within 1 hour, 28 genes showed more than 4-fold increment within 4 hours but not upregulated within 1 hour. We combined those “Late upregulated genes” with 508 genes and referred to as “All upregulated genes” in the subsequent analysis (Figure S1). Mapping the classified genes to central carbon metabolism revealed that *nifJ* encoding PFOR and *hox* operon encoding a bidirectional hydrogenase complex were transiently upregulated (Figure 1D and Table 1). RTqPCR verified the transient expression of *hoxF*, *hoxH,* and *nifJ* (Figure S2).

**Figure 1.**
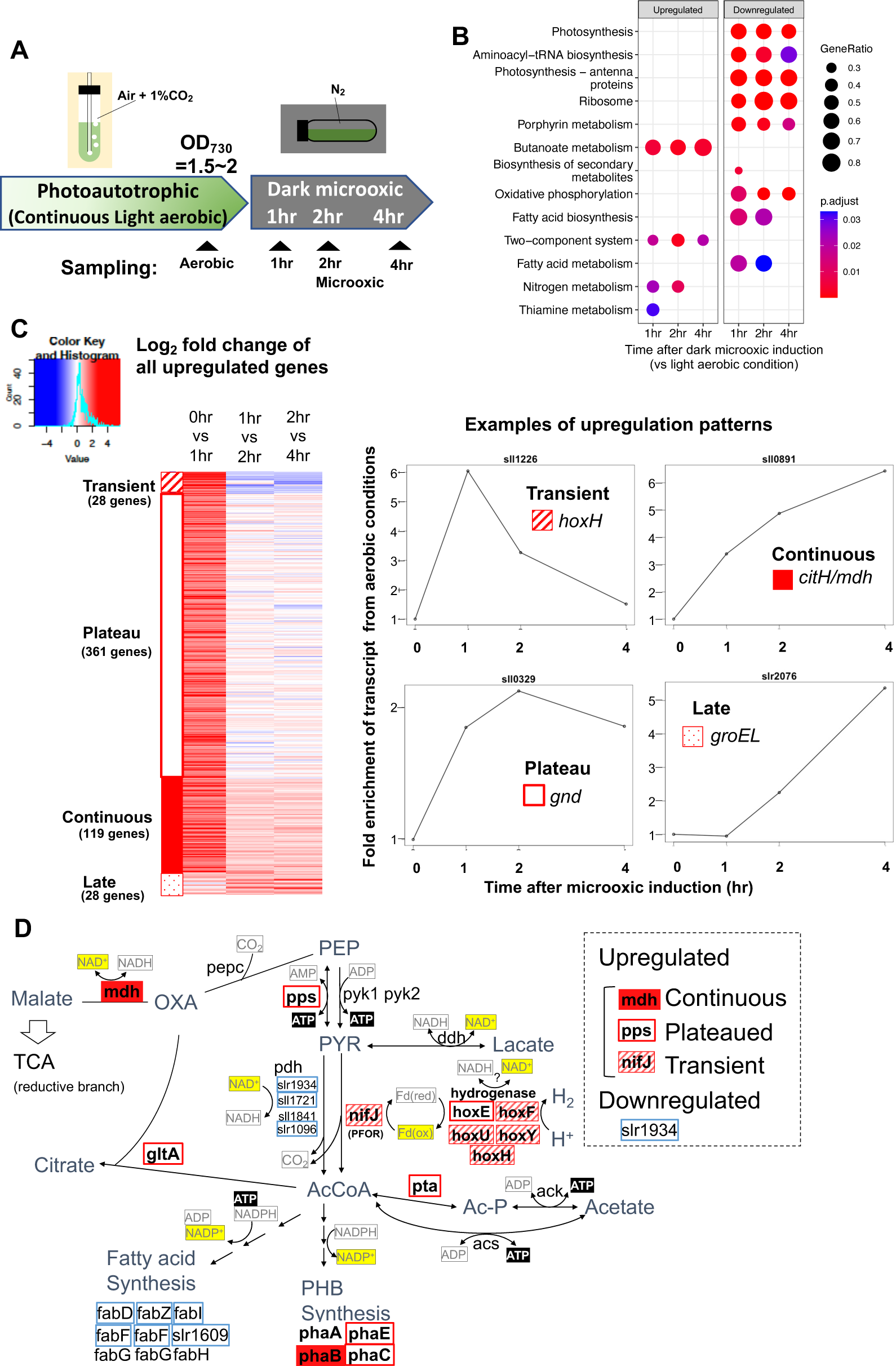
(A) Schematic diagram for the sampling of cells under aerobic and microoxic conditions. (B) Gene set enrichment analysis on time-course transcriptome data. KEGG pathways enriched in upregulated or downregulated genes after 1-, 2-, and 4-h incubation under microoxic conditions are shown. (C) (left) Heatmap showing expression change in all upregulated genes over the time-course. Genes classified into Transient (striped square), Plateau (open square), Continuous (filled square), and Late (dotty square) were clustered into subgroups and sorted by the gene name. (right) Examples of genes are classified into each expression pattern. (D) The classified genes were mapped to central carbon metabolism, centered on pyruvate. PEP: phosphoenolpyruvate, PYR: pyruvate, AcCoA: acetyl CoA, Ac-P: acetyl phosphate, OXA: oxaloacetate, PHB: polyhydroxy butyrate. TCA: tricarboxylic acid cycle.

**Table 1.**
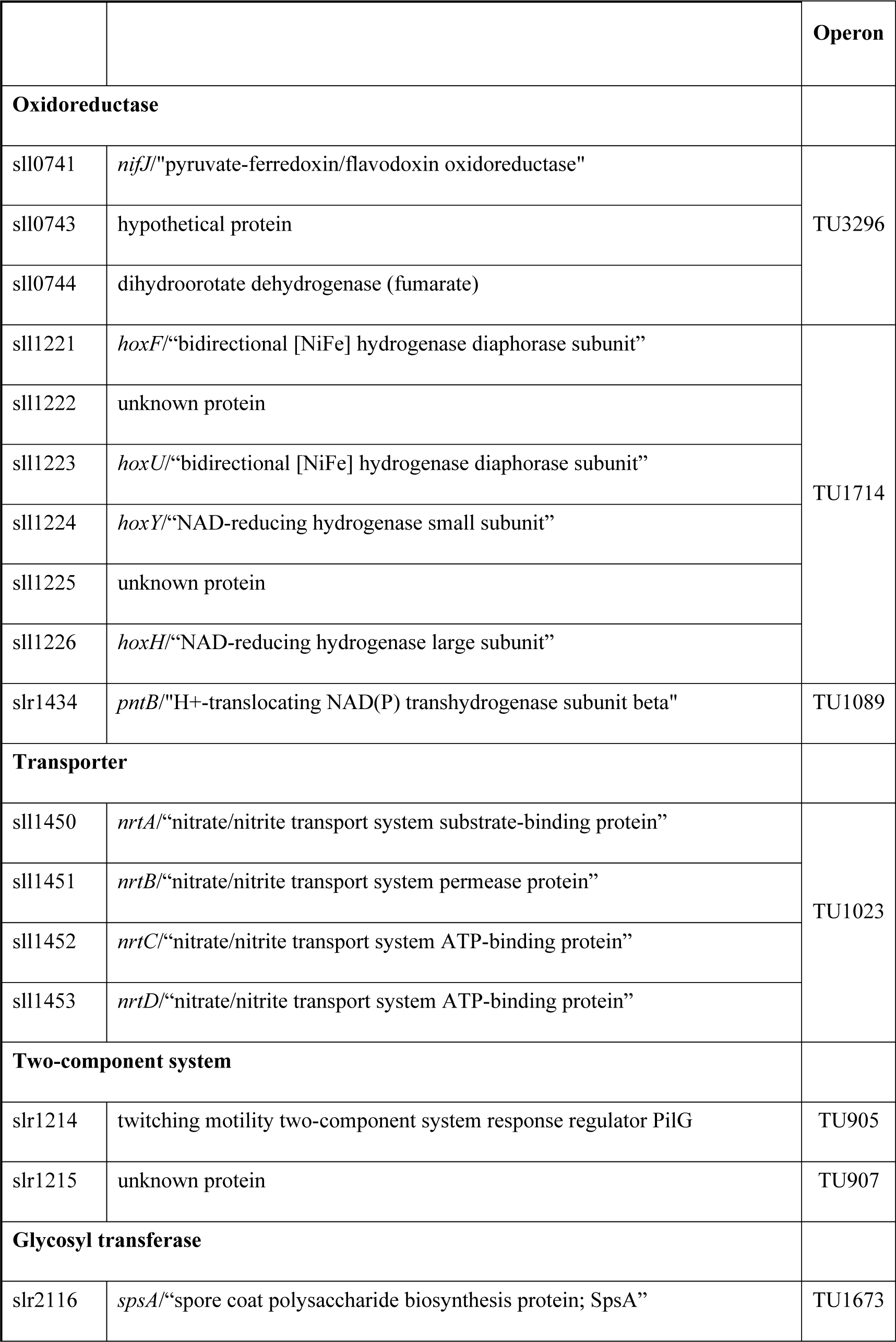
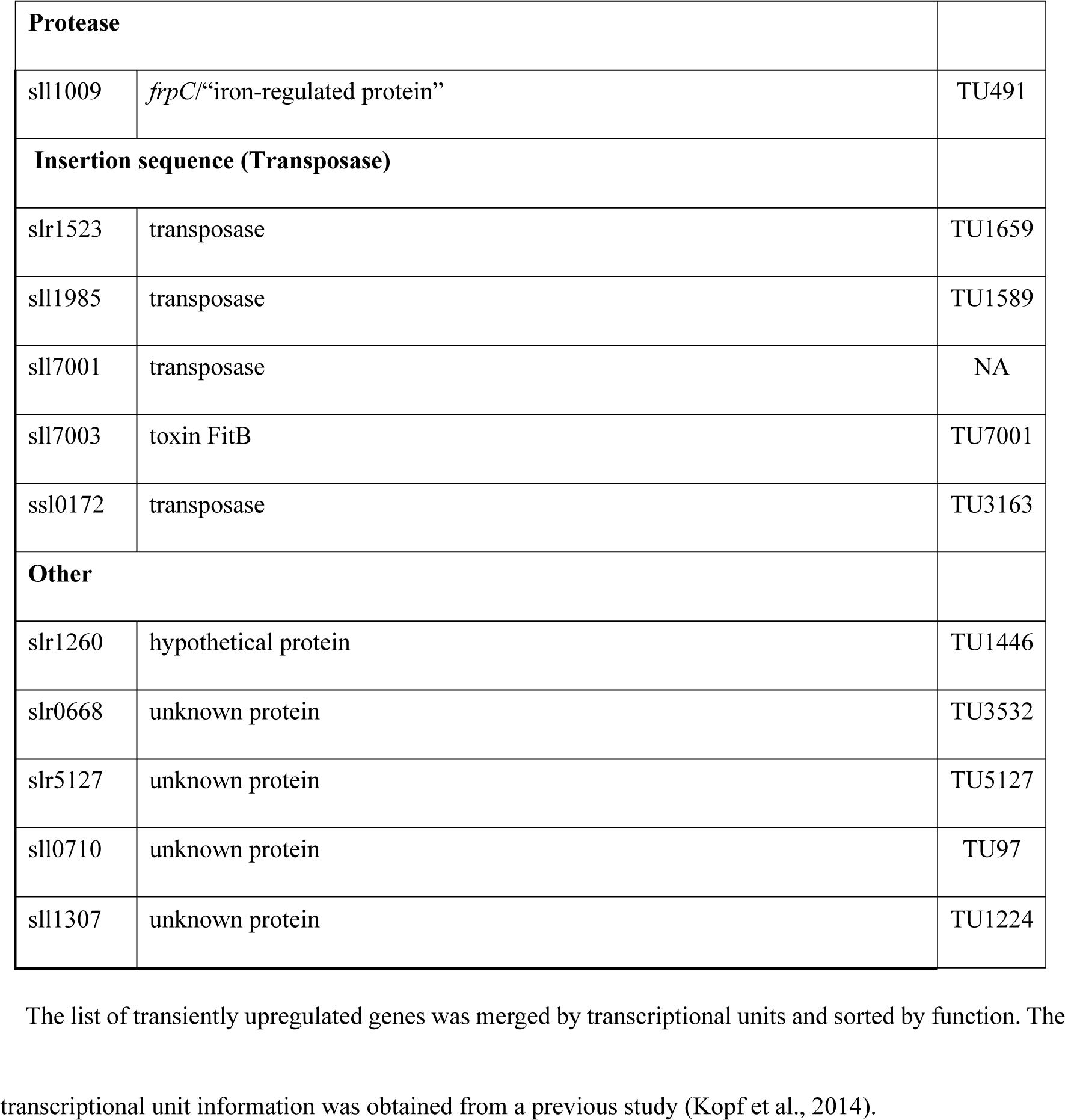
List of transiently upregulated genes.

### SigE and cyAbrB2 control the expression of transiently upregulated genes

The functional correlation between hydrogenase and PFOR, encoded by the *hox* operon and *nifJ,* suggests that transient upregulation has physiological significance. We focused on transiently upregulated genes and attempted to reveal the regulatory mechanism underlying transient upregulation. While SigE promotes the expression of *hox* and *nifJ*, cyAbrB2 represses them (17, 19, 20). We confirmed that the deletion of *sigE* and *cyabrb2* (Δ*sigE* and Δ*cyabrb2*, respectively) affected the expression of *hoxF*, *hoxH*, and *nifJ* by RT-qPCR (Figure S1). Thus, we conducted a time-course transcriptome analysis of Δ*sigE* and Δ*cyabrb2* under aerobic conditions and after 1- and 2-hours cultivation in dark microoxic conditions (Figures 2A and S3). The transcriptome data confirmed that SigE and cyAbrB2 regulated *hox* operon expression (Figure 2B). At each time point, we searched for differentially expressed genes (DEGs) between mutants and wildtype with a more than 2-fold expression change and a false discovery rate (FDR) less than 0.05. We found that deleting *sigE* or *cyabrb2* preferentially affected the expression of transiently upregulated genes, not limited to *hox* and *nifJ* operons (Figures 2C and 2D). Interestingly, *cyabrb2* deletion resulted in the upregulated expression of transient genes under aerobic conditions, in contrast to 1-hour cultivation under microoxic conditions (Figure 2C).

**Figure 2.**
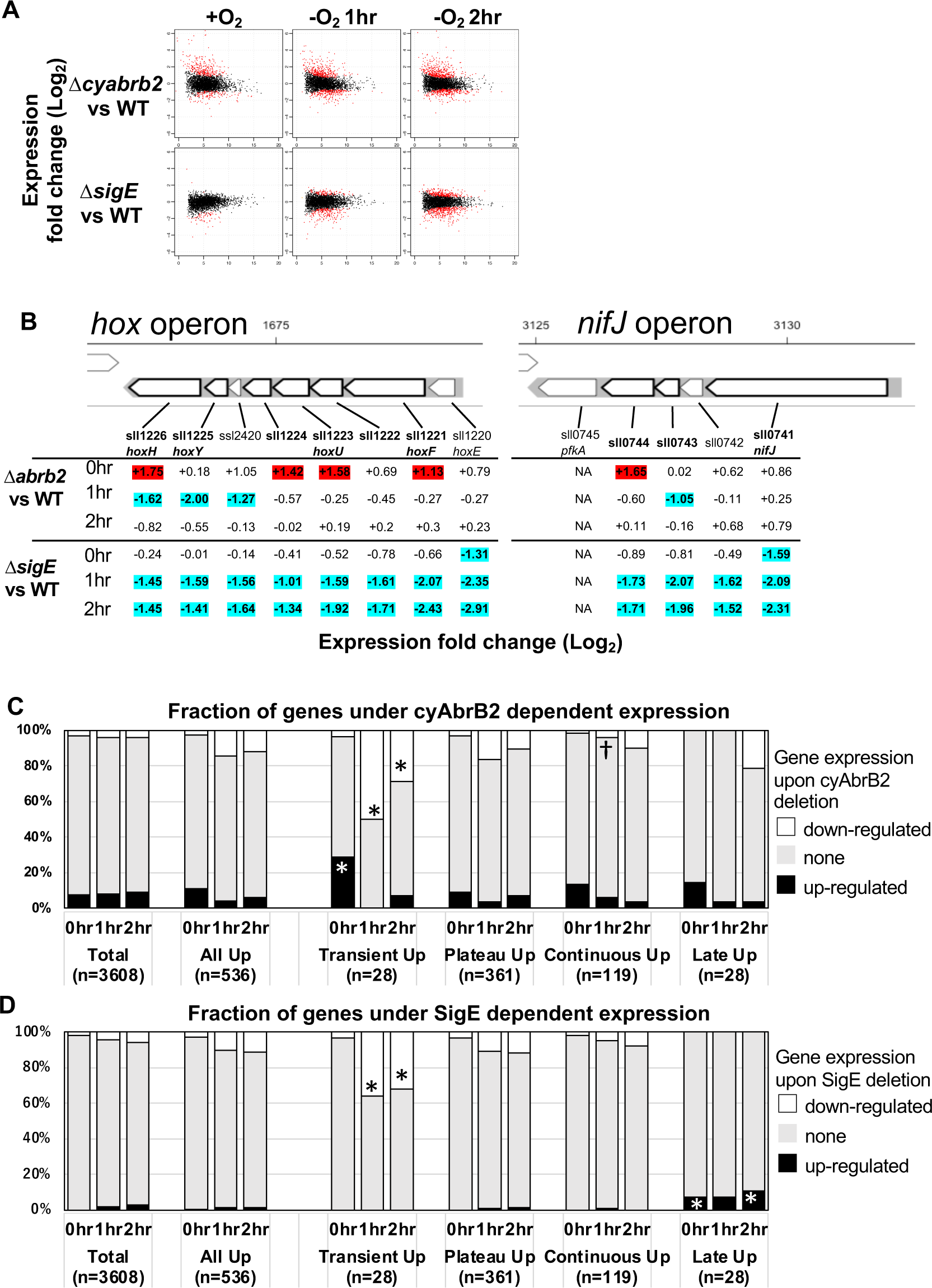
(A) MA plot showing fold change (y-axis) and average (x-axis) of gene expression between wildtype and mutant strains at each timepoint. Red dots indicate defined DEGs (|Log_2_ FC| >1 with FDR < 0.05). (B) Log2 scaled expression fold change in genes in the *hox* and *nifJ* operons upon Δ*cyabrb2* and Δ*sigE* under aerobic conditions (0 h), 1 hour after microoxic condition (1 hr), and 2 hours after microoxic condition (2 hr). DEGs are marked with sky blue (downregulated upon deletion) or red (upregulated upon deletion). (C and D) Fraction of upregulated and downregulated genes upon the (C) Δ*cyabrb2* and (D) Δ*sigE* at the timepoints of aerobic conditions (0 hr), 1 hour after anoxic condition (1 hr), and 2 hour after anoxic condition (2 hr). Genes are classified according to Figure 1C. Asterisk (*) and dagger (†) denote statistically significant enrichment and anti-enrichment compared with all upregulated genes tested by multiple comparisons of Fisher’s exact test (FDR < 0.05).

### Genome-wide analysis of cyAbrB2, cyAbrB1, and SigE via ChIP-seq

To decipher the regulatory mechanism of transiently upregulated genes, we must first comprehend the fundamental features and functions of these transcriptional regulators. Therefore, a genome-wide survey of cyAbrB2 and SigE occupation (Figure S4) combined with transcriptome data was done. Specifically, we generated a *Synechocystis* strain in which cyAbrB2 was epitope-tagged and performed a ChIP-seq assay under aerobic and microoxic conditions (Figure S4A). SigE-tagged strains previously constructed and analyzed elsewhere were also employed (25). The primary sigma factor SigA was also analyzed to determine SigE-specific binding. In addition to cyAbrB2, we tagged and analyzed cyAbrB1, which is the interactor of cyAbrB2 and positively regulates the *hox* operon.

### CyAbrB2 binds to long tracts of the genomic region and suppresses genes in the binding region

The ChIP-seq data showed that cyAbrB2 bound to long tracts of the genomic region with lower GC content than the whole genome *Synechocystis* (Figures 3A and 3B). Vice versa, regions exhibiting lower GC contents showed a greater binding signal of cyAbrB2 (Figure 3C). This correlation was not a systematic bias of next-generation sequencing because the binding signals of SigE, SigA, and control showed no negative correlation to GC contents (Figures S5A and S5B). The binding regions of cyAbrB2 called by peak-caller covered 15.7% of the entire genome length. 805 of 3614 genes overlapped with cyAbrB2 binding regions, and almost half (399 of 805 genes) were entirely covered by cyAbrB2 binding regions. The cyAbrB2 binding regions included 80 of 125 insertion sequence elements (Figure 3D). Comparison with the transcriptome of Δ*cyabrB2* revealed that cyAbrB2 tended to suppress the genes overlapping with its binding regions under aerobic conditions (Figures 3A and 3E). A survey of the genomic localization of cyAbrB1 under aerobic conditions revealed that cyAbrB1 and cyAbrB2 shared similar binding patterns (Figures 3A and S5C). Due to the essentiality of cyAbrB1, we did not perform transcriptome analysis on a cyAbrB1-disrupted strain. Instead, we referred to the recent study performing transcriptome of cyAbrB1 knockdown strain, whose cyAbrB1 protein amount drops by half (30). Among 24 genes induced by cyAbrB1 knockdown, 12 genes are differentially downregulated genes in *cyabrb2*Δ in our study (Figure S5D).

**Figure 3.**
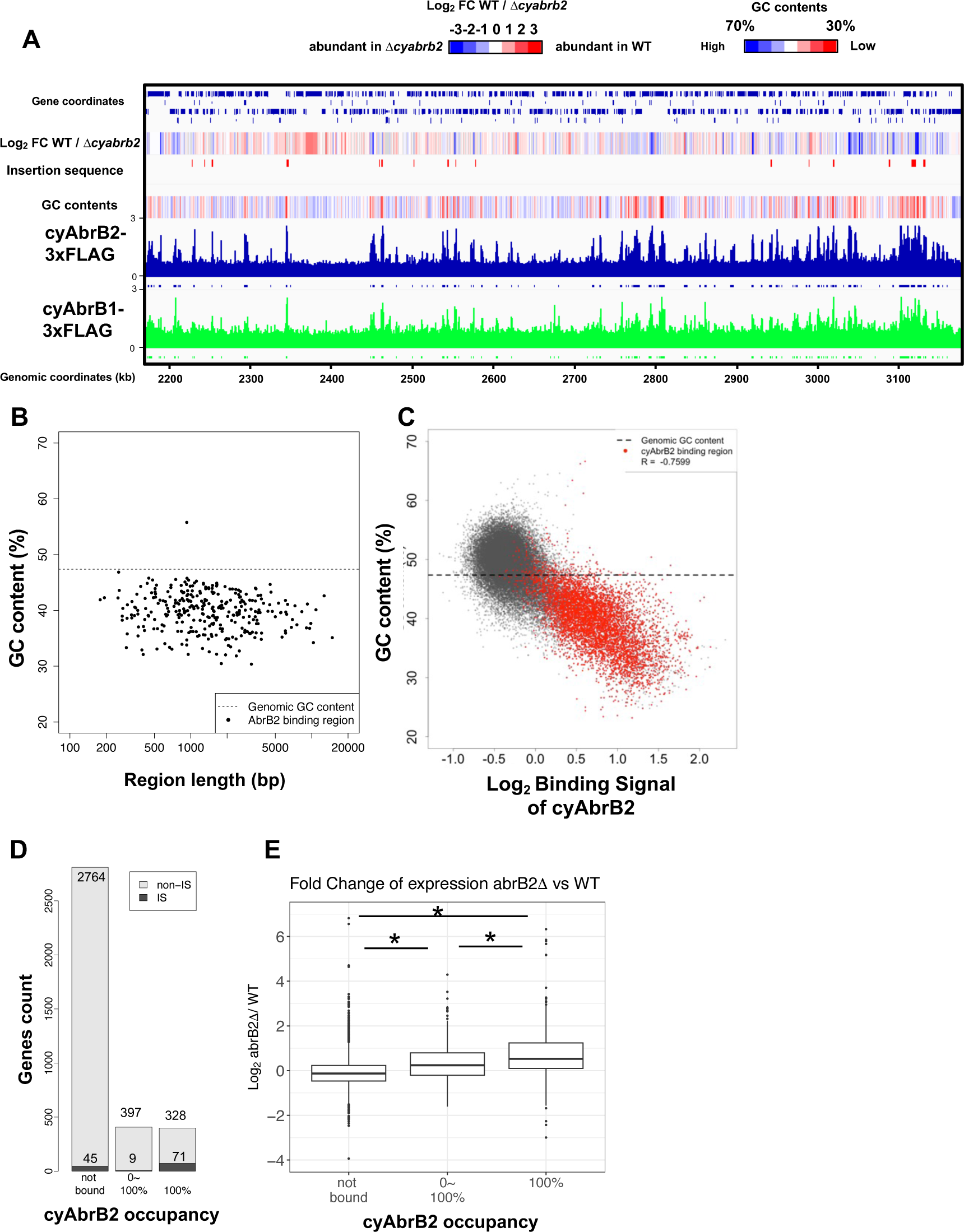
(A) Snapshot of ChIP-seq data for cyAbrB2 and cyAbrB1 under aerobic conditions. The heatmap in the second column indicates expression fold change upon Δ*cyabrb2* under aerobic conditions. Positive values (colored in red) indicate that the gene expression is higher in wildtype than in Δ*cyabrb2*, and negative values (colored in blue) indicate the opposite. The positions for the insertion elements are marked with red in the third column. The heatmap in the fourth column indicates GC contents. High GC contents are colored in blue and low GC contents are colored in blue. (B) GC contents and region length of cyAbrB2 binding regions (black dots). The horizontal dotted line indicates the genomic average of GC content. (C) Scatter plot of GC content and binding signal of cyAbrB2. The x-axis is the binding signal of cyAbrB2 in each 100 bp region, and the y-axis indicates GC contents within 500 bp windows sliding every 100 base pairs. CyAbrB2 binding regions are marked with red dots. (D) Histogram of genes showing the extent of occupancy (not bound, partially overlapped, or entirely overlapped) by the cyAbrB2 binding region. The gray bars indicate non-IS genes, and the count numbers of the non-IS genes are displayed on the gray bars. The black bars indicate the IS genes, and the count numbers of the IS genes are displayed above the black bars. (E) Boxplot showing fold change in gene expression by Δ*cyabrb2* under aerobic conditions. Genes are classified according to the extent of occupancy by the cyAbrB2 binding region. Asterisk (*) denotes statistical significance tested by multiple comparisons of the Wilcoxon-rank test.

### CyAbrB2 binds to transiently upregulated genes

The binding regions of cyAbrB2 overlapped 17 of 28 transiently upregulated genes, showing significant enrichment from all upregulated genes (Figure 4A). The transiently upregulated genes belong to 17 transcriptional units (TUs), according to the previous study(31), and cyAbrB2 tends to bind TUs with transiently upregulated genes (Figure 4B). While cyAbrB2 covered the entire length of insertion sequences and unknown proteins, its binding positions on other transient genes were diverse (Figure 4C). Specifically, the *hox* and *nifJ* operons had two distinct binding regions located at the transcription start sites (TSSs) and middle of operons, the *pntAB* operon had two binding regions in the middle and downstream of the operon, and the *nrtABCD* operon had one binding region downstream of the operon (Figure 4C).

**Figure 4.**
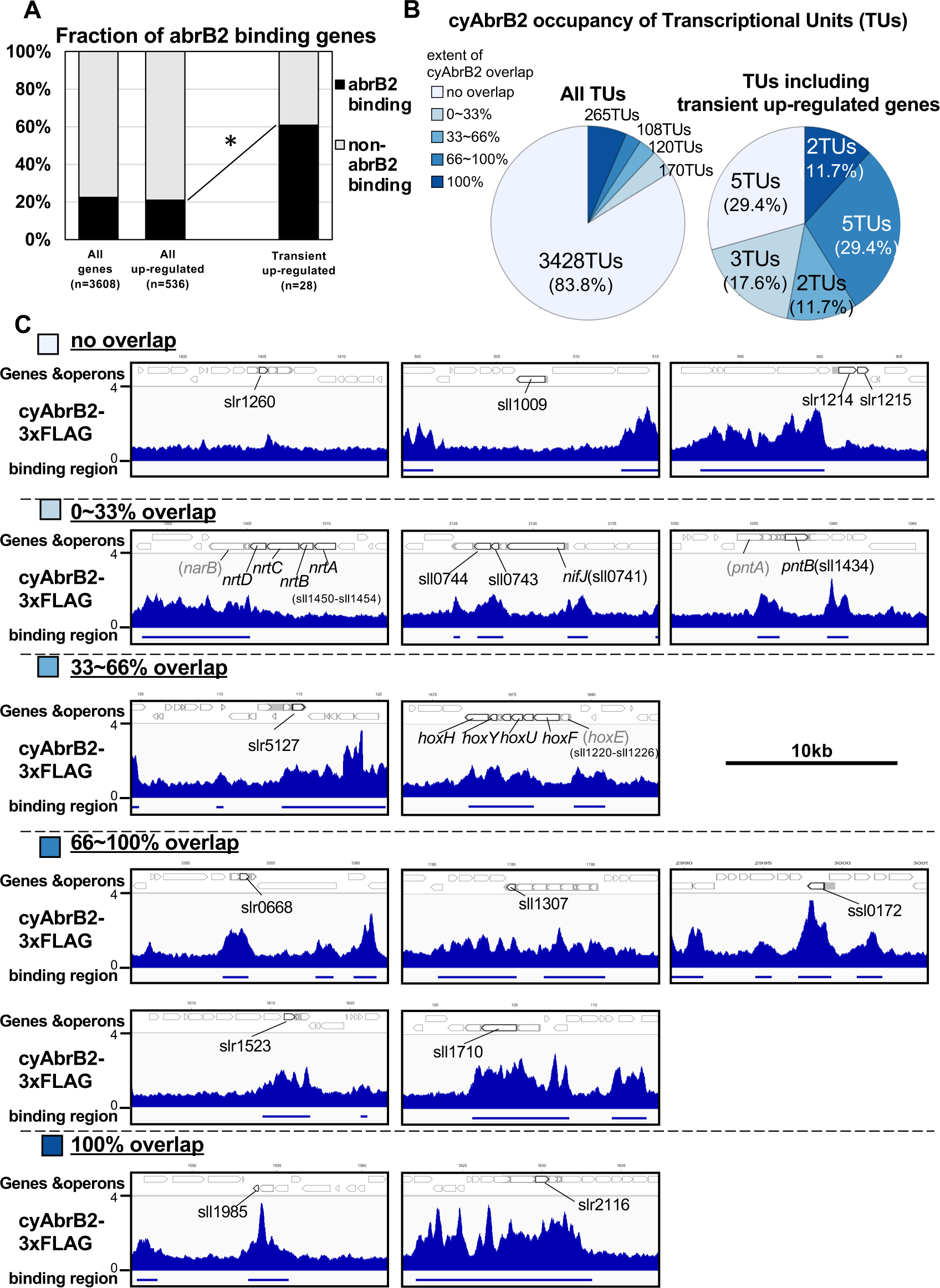
(A) Fraction of genes overlapped or non-overlapped with cyAbrB2 binding regions at the timepoints of aerobic conditions. Genes are classified according to Figure S1. Asterisk (*) denotes statistically significant enrichment compared with all upregulated genes tested by multiple comparisons of Fisher’s exact test. (B) Pie charts of transcriptional units (TUs) classified by extent of overlapping with cyAbrB2 binding region. The left pie represents all TUs, and the right pie represents only TUs containing the transient upregulated genes. (C) Distribution of cyAbrB2 in the aerobic condition around transiently upregulated genes. Arrows with bold lines indicate transiently upregulated genes. Shaded arrows indicate operons whose data were obtained from a previous study. The bars below the graph indicate the binding regions of each protein. The black bar at the top of the figure indicates a length of 10 kbp.

### Localization of cyAbrB2 became blurry under the microoxic condition

When cells entered microoxic conditions, the relative ChIP-seq signals in the cyAbrB2 binding regions slightly declined (Figures 5A and 5B). Notably, the total quantities of precipitated DNA by tagged cyAbrB2 did not decrease (Figure S6A), and qPCR confirmed that the cyAbrB2 binding signal increased in all positions tested (Figure 5C). ChIP-seq data and ChIP-qPCR data indicate that the boundary between cyAbrB2 binding region and cyAbrB2-free region became obscured when the cells entered microoxic conditions due to increased binding of cyAbrB2 to both cyAbrB2 binding and cyAbrB2-free region. The protein amount of cyAbrB2 was not altered on entry to the microoxic condition (Figure S6B). The cyAbrB2 binding signal around the transiently upregulated genes became less specific upon entry into microoxic conditions, consistent with the general tendency (Figure 5B). The amount of DNA immunoprecipitated by cyAbrB1 was also increased in the microoxic condition, and the protein amount was not increased (Figures S6C and S6D).

**Figure 5.**
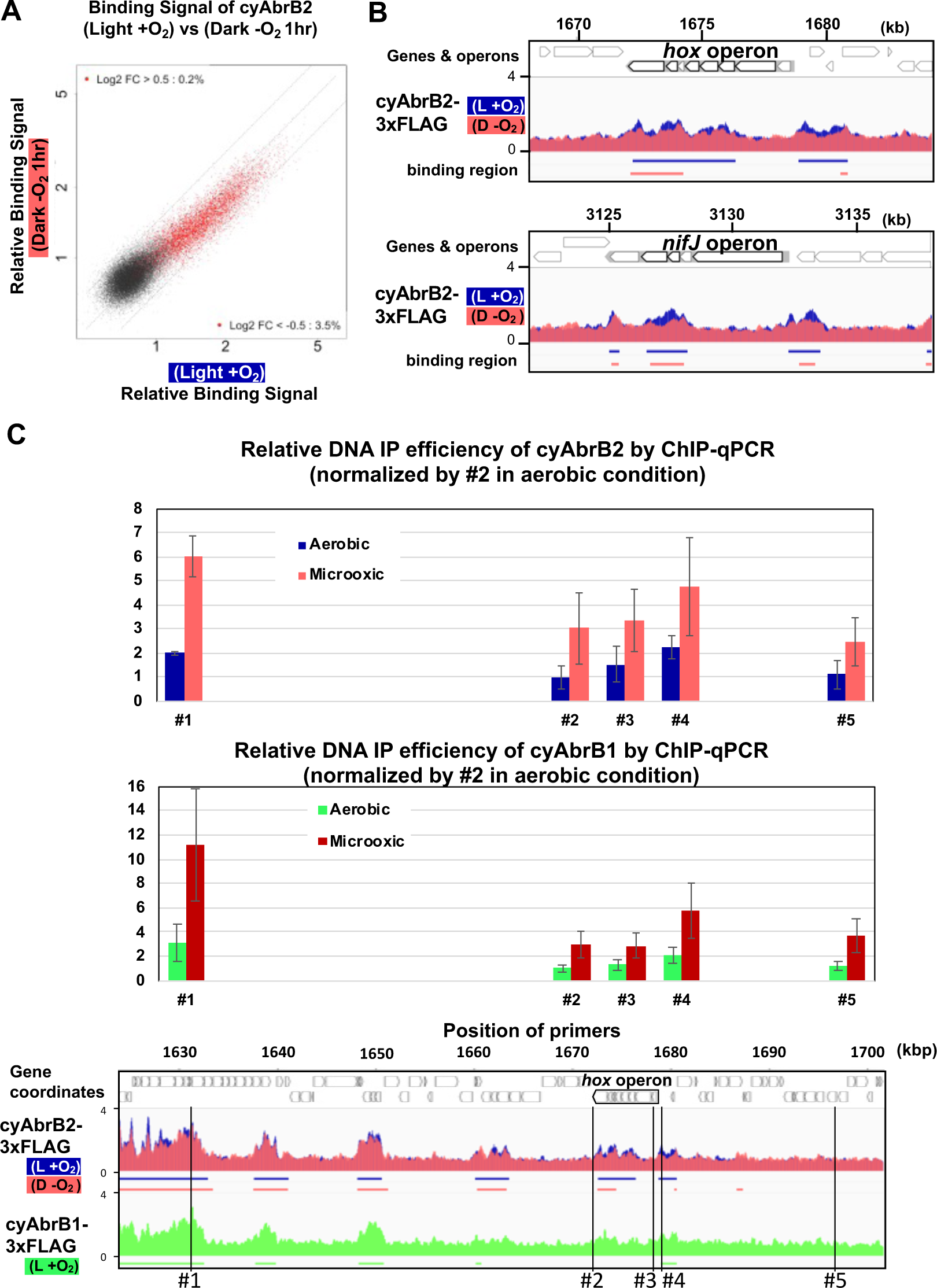
(A) Scatter plot showing changes of the binding signal by 1hr cultivation in the microoxic condition. The binding signal of each 100bp window is plotted. Red dots are cyAbrB2 binding regions in either aerobic or microoxic conditions. The dotty lines indicate Log2 fold enrichment of 0.5, 0, and −0.5 between aerobic and microoxic conditions. (B) Distribution of cyAbrB2 around *hox* operon and *nifJ* operon. ChIP-seq data in aerobic (L + O_2_) and dark microoxic (D − O_2_) conditions are overlayed. The bars below the graph indicate the binding regions of each protein. (C) Quantification for IP efficiency of cyAbrB2 (top) and cyAbrB1 (middle) by qPCR in the aerobic and microoxic conditions. The position of primers and ChIP-seq data of cyAbrB2 are shown at the bottom. Scores are normalized by the IP% at position #2 in the aerobic condition.

### Sigma factors SigE and SigA are excluded from cyAbrB2 binding regions regardless of GC contents

We searched for SigE and SigA binding sites under aerobic and microoxic conditions (Figures S7A and S7B, left and right, respectively). The SigE and SigA peaks identified in this study predominantly covered the previously identified peaks (Figure S7C), reproducing the previous study’s conclusion (25); i.e., SigE and the primary sigma factor SigA share localization on the promoters of housekeeping genes, but SigE exclusively binds to the promoters of its dependent genes. SigE and SigA binding peaks were significantly excluded from the cyAbrB2 binding regions (Figures 6A and 6B). SigE preferred the cyAbrB2-free region in the aerobic condition more than SigA did (Odds ratios of SigE and SigA being in the cyAbrB2-free region were 4.88 and 2.74, respectively). CyAbrB2 prefers AT-rich regions, but no correlation was found between the GC content and binding signals of SigE and SigA (Figures S5A and S5B). Thus, SigA and SigE are excluded from cyAbrB2 binding regions regardless of GC contents.

**Figure 6.**
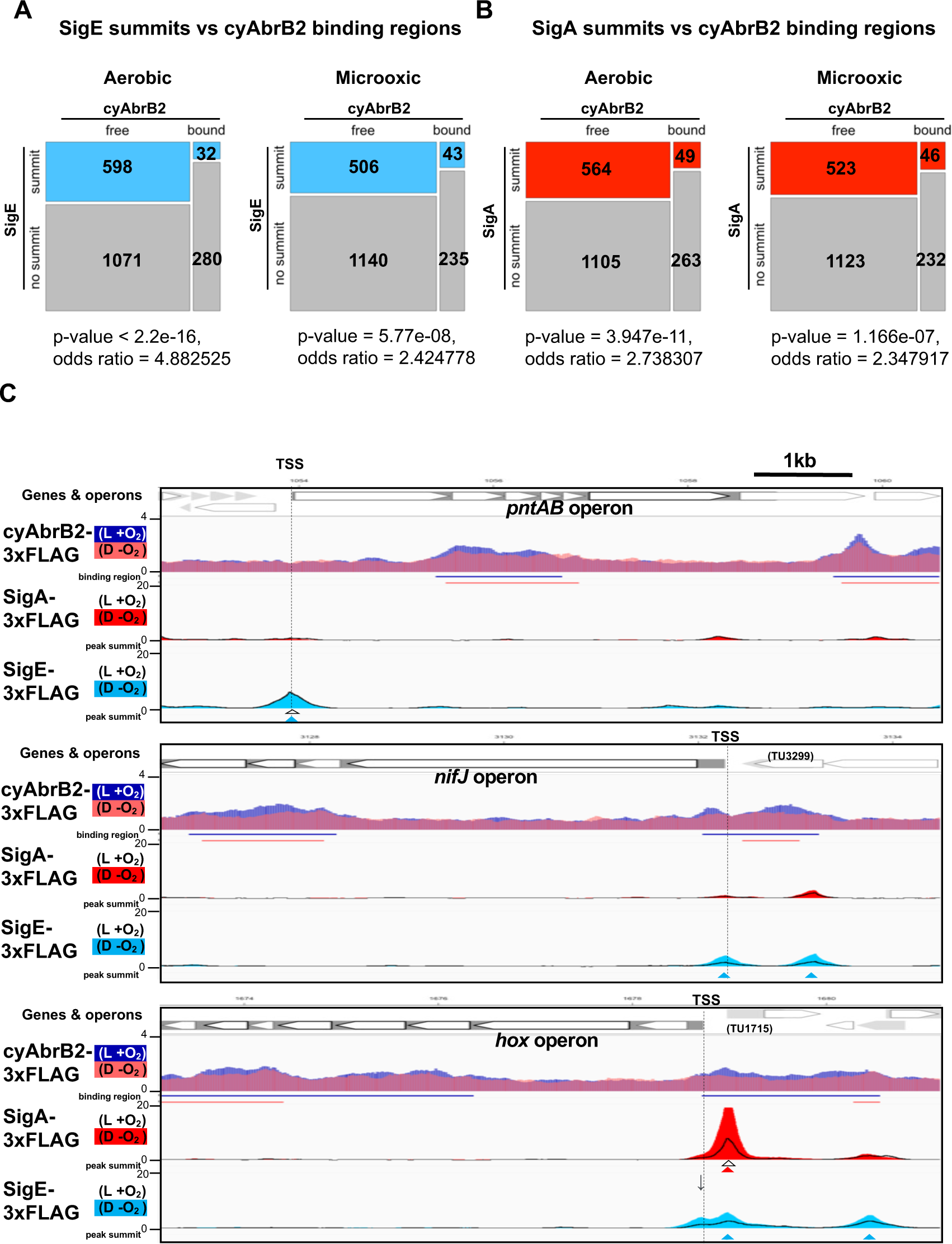
(A, B) Anti-co-occurrence of cyAbrB2 binding regions and sigma factors. Mosaic plots of cyAbrB2 binding regions and SigE peaks (A) or SigA binding peaks (B) are shown. Odds and p-values were calculated by Fisher’s exact test. (C) Snapshots of ChIP-seq data for CyAabrB2, SigE, and SigA at the *nifJ* region (top) and *hox* region (bottom). ChIP-seq data for cyAbrB2, SigE, and SigA under aerobic and dark microoxic conditions are overlayed. ChIP-seq data of cyAbrB2 under aerobic and microoxic conditions are colored blue and pink, respectively. ChIP-seq data for SigE and SigA are shown in solid lines (aerobic conditions) and the area charts (microoxic conditions). The positions of TSSs were obtained from a previous study (Kopf et al., 2014) and indicated by vertical dotted lines. Open triangles indicate peak summits under aerobic conditions, and solid triangles indicate peak summits under microoxic conditions.

### Dynamics of sigma factors upon exposure to the microoxic condition

When cells entered microoxic conditions, the binding signals of SigA and SigE were changed, although most of their peaks observed under aerobic conditions were present under microoxic conditions (Figure S7A). The preference of SigE for the cyAbrB2-free region was alleviated in the microoxic condition (Figure 6A). Next, we focused on sigma factor dynamics in transiently upregulated genes. SigE, but not SigA, binds at the TSS of *pntAB* under aerobic and microoxic conditions (Figure 6C top). SigE binding summits were not identified at the TSSs of the *hox* and *nifJ* operons under aerobic conditions. However, the SigE-specific binding summit appeared at the TSS of *nifJ* when cells entered microoxic conditions (Figure 6C middle). A bimodal peak of SigE was observed at the TSS of the *hox* operon in a microoxic-specific manner (Figure 6C bottom panel). The downstream side of the bimodal SigE peak coincides with the SigA peak and the TSS of TU1715. Another side of the bimodal peak lacked SigA binding and was located at the TSS of the *hox* operon (marked with an arrow in Figure 6C), although the peak caller failed to recognize it as a peak. SigE binding without SigA on the promoters of *hox*, *nifj*, and *pntAB* is consistent with their SigE-dependent expression (Figure 2B).

### Chromatin conformation around *hox* operon and *nifJ* operon

We have shown that cyAbrB2 broadly binds to AT-rich genomic regions, including insertion element sequences, and represses expression (Figure 3). This is functionally similar to the NAPs (14), which makes us hypothesize that cyAbrB2 modulates chromosomal conformation. Therefore, we conducted the chromatin conformation capture (3C) assay against wildtype and *cyabrb2*Δ strains at aerobic and microoxic conditions. qPCR was performed with unidirectional primer sets, where the genomic fragment containing *hox* operon and *nif* operon (hereinafter *hox* fragment and *nifJ* fragment, respectively) were used as bait (Figure 7).

**Figure 7.**
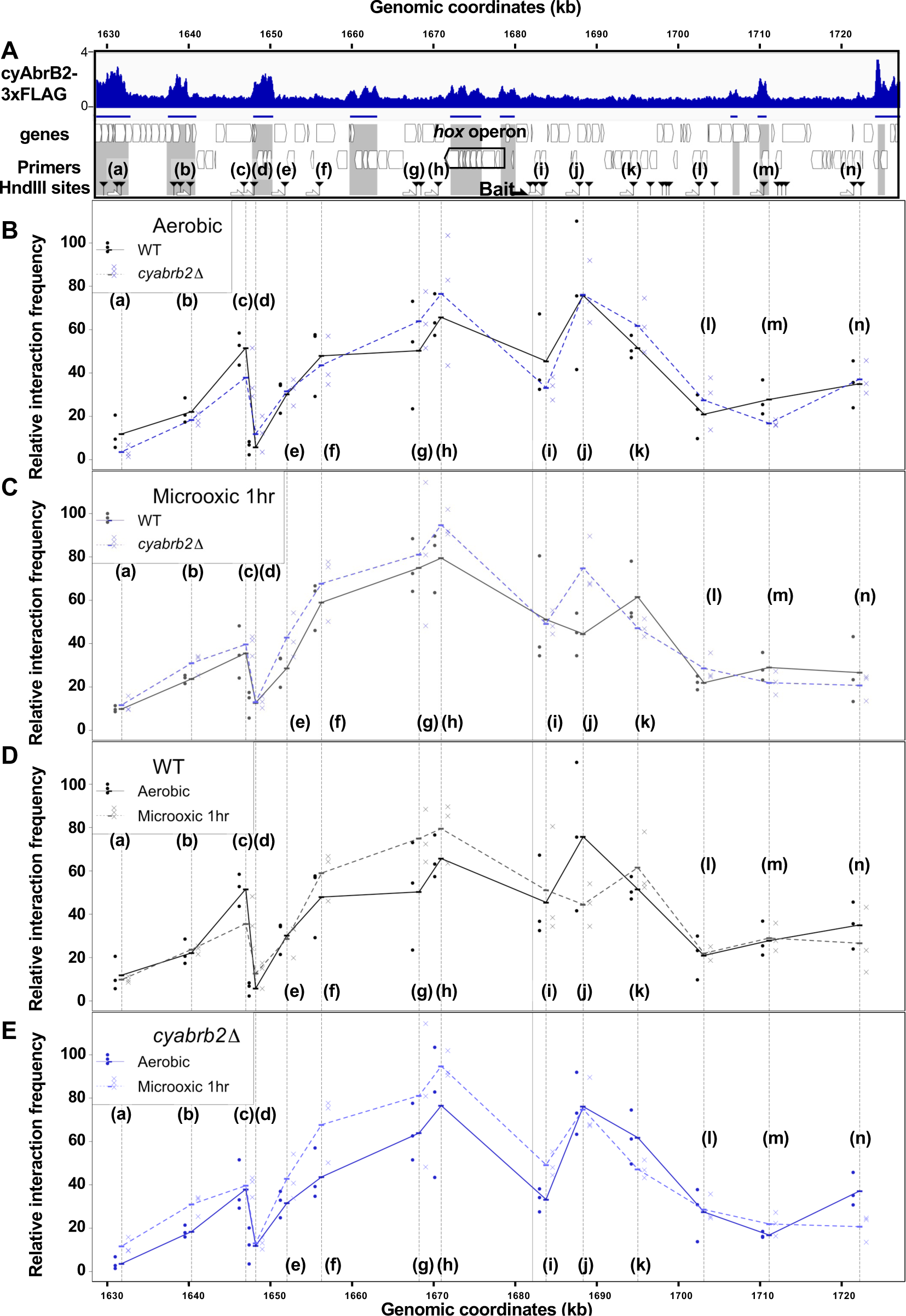
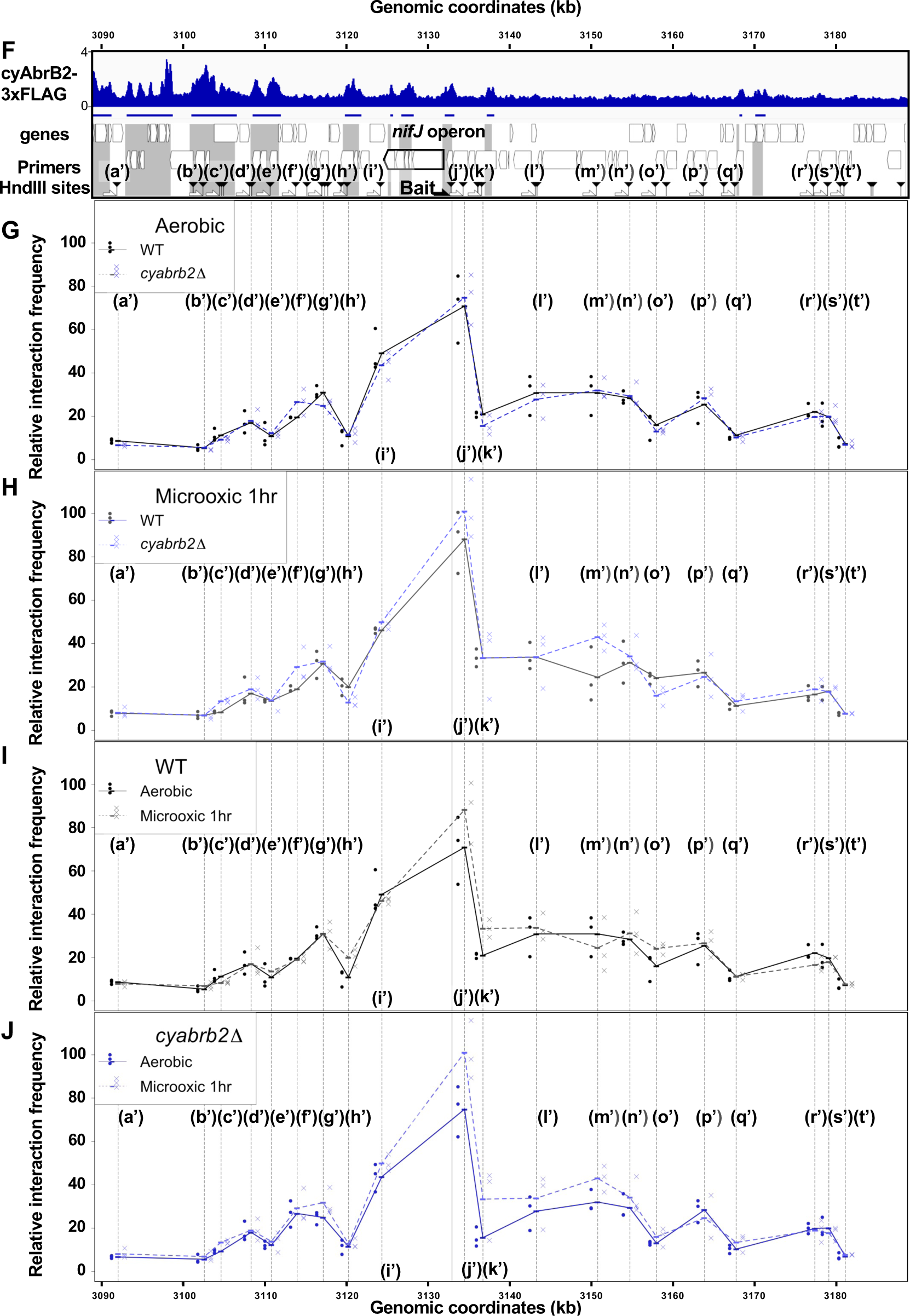
(A) and (F)Schematic diagram of 3C analysis around *hox* operon (A) and *nifJ* operon (F). In the panels (A) and (F), the black horizontal arrow shows the location of the bait primer, and white horizontal arrows ((a) to (n) in *hox* operon (A) and (a’) to (t’) in *nifJ* operon (F)) indicate loci where the interaction frequency with bait were assayed. Vertical black arrowheads indicate the position of HindIII sites. ChIP-seq data of cyAbrB2 in the aerobic condition is displayed in the bottom, and cyAbrB2 binding regions are marked with shade. (B - E) The line plot showing the interaction frequency of each locus with *hox* fragment. Two of data sets are presented; (B) wildtype vs Δ*cyabrb2* in aerobic condition, (C) wildtype vs Δ*cyabrb2* in 1hr of microoxic condition, (E) wildtype in aerobic vs 1hr of microoxic condition, and (E) Δ*cyabrb2* in aerobic vs 1hr of microoxic condition are compaired. (G – J) The line plot showing the interaction frequency of each locus with *nifJ* fragment. Two data sets are selected and presented; (G) wildtype vs Δ*cyabrb2* in aerobic condition, (H) wildtype vs Δ*cyabrb2* in 1hr of microoxic condition, (I) wildtype in aerobic vs 1hr of microoxic condition, and (J) Δ*cyabrb2* in aerobic vs 1hr of microoxic conditions are compared. The line plots indicate the average interaction frequency over the random ligation (n=3). Individual data are plotted as dots.

First, focusing on the aerobic condition of wildtype (Figure 7B, solid line), the *hox* fragment interacted with its proximal downstream loci (loci (f) to (g)) and proximal upstream locus (locus (j)). The *hox* fragment also interacts with the distal downstream locus (locus (c)). Meanwhile, the *nifJ* fragment shows high interaction frequency with proximal upstream and downstream loci (Figure 7G, loci (i’) and (j’)), and a distal downstream locus (locus (g’)) showed higher interaction frequency with *nifJ* fragment than proximal locus (h’) did. The upstream regions of *nifJ* (loci (l’) to (n’) and (p’)) showed comparable frequency with locus (g’).

### The chromatin conformation is changed in *cyabrb2*Δ in some loci

Then we compared the chromatin conformation of wildtype and *cyabrb2*Δ. Although overall shapes of graphs did not differ, some differences were observed in wildtype and *cyabrb2*Δ (Figures 7B and 7G); interaction of locus (c) with *hox* region were slightly lower in *cyabrb2*Δ and interaction of loci (f’) and (g’) with *nifJ* region were different in wildtype and *cyabrb2*Δ. Note that the interaction scores exhibit considerable variability and we could not detect statistical significance at those loci.

### Changes of chromatin conformation upon microoxic condition

When the cells entered the microoxic condition, proximal loci interacted more frequently (Figure 7D; loci (f)-(h) and Figure 7I loci (j’) and (k’)). This tendency was more apparent in *cyabrb2*Δ (Figures 7E and 7J). Furthermore, the interaction of *nifJ* upstream loci (l’) – (n’) increased in the microoxic condition in *cyabrb2*Δ but not wildtype (Figures 7I and 7J). The locus (c) and locus (j) interacted less frequently with *hox* fragment upon entry to the microoxic condition in the wildtype. While the interaction scores exhibit considerable variability, the individual data over time demonstrate declining trends of the wildtype at locus (c) and (j) (Figure S8). In Δ*cyabrb2*, by contrast, the interaction frequency of loci (c) and (j) was unchanged in the aerobic and microoxic conditions (Figure 7E). The interaction frequency of locus (c) in Δ*cyabrb2* was as low as that in the microoxic condition of wildtype, while that of locus (j) in Δ*cyabrb2* was as high as that in the aerobic condition of wildtype (Figures 7B and 7C). In summary, 3C analysis demonstrated cyAbrB2-dependent and independent dynamics of chromosomal conformation around the *hox* and *nifJ* operon in response to the microoxic condition (Figure 8).

**Figure 8.**
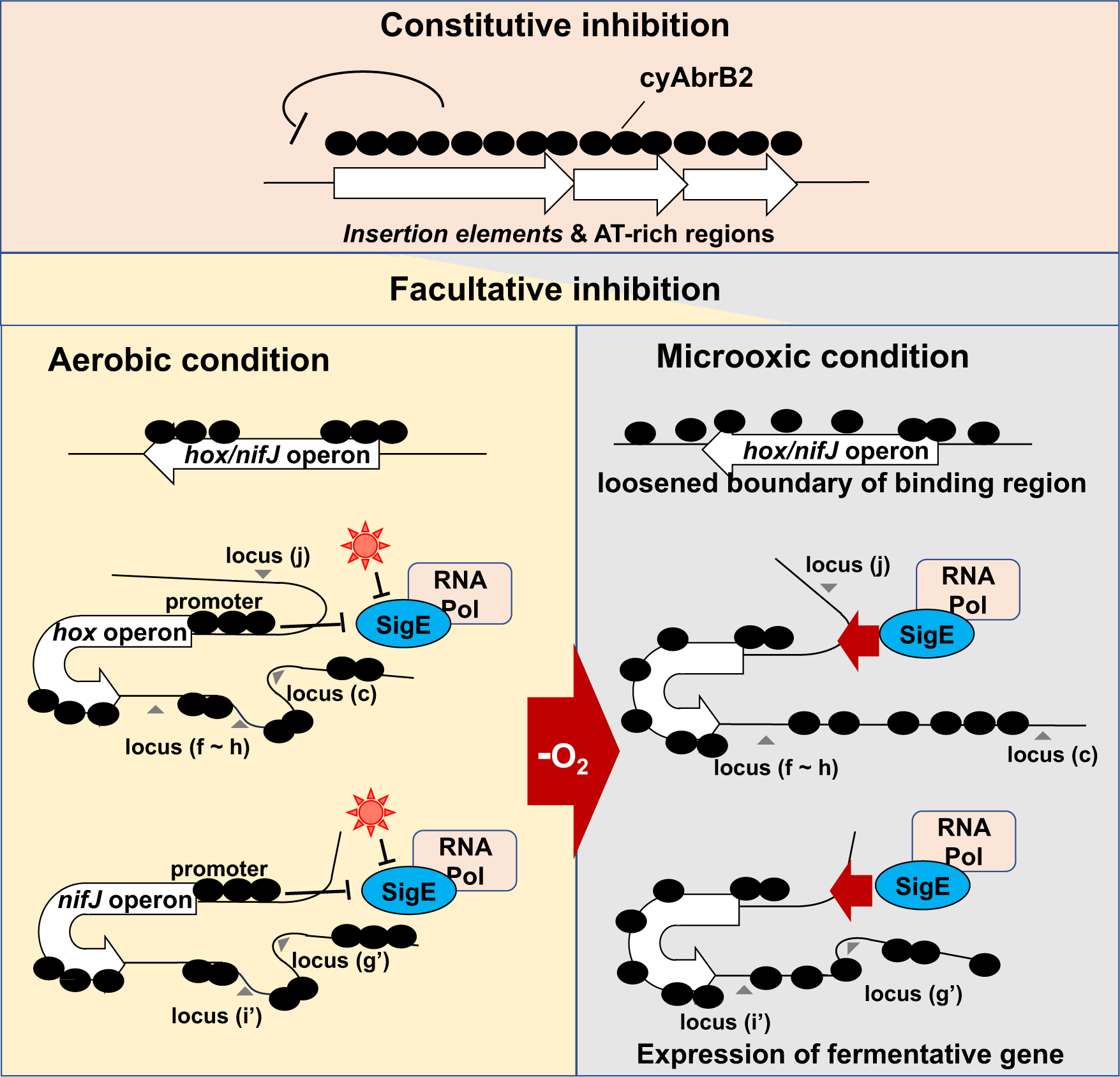
Schematic diagram of the dynamics of transcription factors governing fermentative gene expression.

## Discussion

### Physiological significance of transient upregulation of *hox* and *nifJ* operons

As the transcriptional change can alter the metabolic flow, the transcriptional upregulation of fermentative genes in response to the microoxic condition is expected to be adaptive for energy acquisition and the maintenance of redox balance. Our time-course transcriptome showed upregulation of several genes involved in catabolism upon exposure to the microoxic condition. The transient upregulation of *hox* and *nifJ* operons is distinctive among them (Figure 1D).

One reason for transient upregulation is probably the resource constraints of inorganic cofactors. Hydrogenase and PFOR (the product of *nifJ* gene) have iron-sulfur clusters, and hydrogenase requires nickel for its activity (32, 33). Overexpression of the *hox* operon should be futile under physiological conditions without an adequate nickel supply (34).

Another significance for transient upregulation may be the reusability of fermentative products. Hydrogen, lactate, and dicarboxylic acids can be reused as the source of reducing power when cells return to aerobic conditions (12, 35, 36). The substrate proton is abundant, but hydrogen is diffusive and difficult to store. Therefore, hydrogenase may favor fermentation initiation, and the reductive branch of TCA-producing dicarboxylic acids may become active subsequently. In fact, *citH*/*mdh* (sll0891) encoding a key enzyme of the reductive branch of TCA was classified as continuously upregulated genes in this study (Figures 1C and 1D).

### Mechanisms for transient expression mediated by SigE and cyAbrB2

SigE and cyAbrB2 can independently contribute to the transient transcriptional upregulation. This is evident as the single mutants, Δ*sigE* or Δ*cyabrb2*, maintained transient expression of *hoxF* and *nifJ* (Figure S2). We first discuss cyAbrB2 as the potential NAPs, and then the mechanism of transient upregulation mediated by cyAbrB2 and SigE will be discussed.

### cyAbrB2 is a novel nucleoid-associated protein of cyanobacteria

We have shown that cyAbrB2 broadly binds to AT-rich genomic regions, including IS elements (Figure 3). This is functionally similar to the histone-like nucleoid protein H-NS family; including H-NS in *Enterobacteriaceae* (37, 38)*, and* Lsr2 in *Mycobacteria* (39). Like H-NS and Lsr2, cyAbrB2 may defend against exogenous DNA elements, which often have different GC content. Interestingly, Lsr2 controls genes responding to hypoxia, showing a functional analogy with cyAbrB2 (40).

### The biochemistry of cyAbrB2 will shed light on the regulation of chromatin conformation in the future

H-NS proteins often cause bound DNA to bend, stiffen, and/or bridge (14). DNA-bound cyAbrB2 is expected to oligomerize, based on the long tract of cyAbrB2 binding region in our ChIP-seq data. However, no biochemical data mentioned the DNA deforming function or oligomerization of cyAbrB2 in the previous studies, and preference for AT-rich DNA is not fully demonstrated in vitro (19, 27, 41). Moreover, our 3C data did not support bridging at least in *hox* region and *nifJ* region, as the high interaction locus and cyAbrB2 binding region did not seem to correlate (Figure 7). Therefore, direct observation of the DNA-cyAbrB2 complex by atomic force microscopy is the solution in the future.

Not only DNA structural change but also the effect of the post-translational modification can be investigated by biochemistry. The previous studies report that cyAbrB2 is subject to phosphorylation and glutathionylation (42, 43), and pH and redox state alters cyAbrB1’s affinity to DNA (41). Those modifications might respond to environmental changes and be involved in transient expression. Overall, the biochemistry integrating assay conditions (PTM, buffer condition, and cooperation with cyAbrB1) and output (DNA binding, oligomerization, and DNA structure) will deepen the understanding of cyAbrB2 as cyanobacterial NAPs.

### Cooperative and antagonistic function of cyAbrB1 and cyAbrB2

CyAbrB1, the homolog of cyAbrB2, may cooperatively work, as cyAbrB1 directly interacts with cyAbrB2 (29), their distribution is similar, and they partially share their target genes for suppression (Figures 3A S5C and S5D). The possibility of cooperation would be examined by the electrophoretic mobility shift assay of cyAbrB1 and cyAbrB2 as a complex. Despite their similar repressive function, cyAbrB1 and cyAbrB2 regulate *hox* expression in opposite directions, and their mechanism remains elusive. The stoichiometry of cyAbrB1 and cyAbrB2 bound to DNA fluctuates in response to the environmental changes (28), but there was no difference in the behavior of cyAbrB1 and cyAbrB2 around the *hox* region on entry to the microoxic condition.

### Localization pattern and function of cyAbrB2

Herein, we classified three types of binding patterns for cyAbrB2. The first is that cyAbrB2 binds a long DNA tract covering the entire gene or operon, represented by the insertion sequence elements. CyAbrB2 suppresses expression in this pattern (Figure 3E). In the second pattern, cyAbrB2 binds on promoter regions, such as *hox* operon and *nifJ*. The binding on those promoters fluctuates in response to environmental changes, thus regulating expression. This pattern also applies to the promoter of *sbtA* (Na^+^/HCO_3_^−^ symporter), where cyAbrB2 is bound in a CO_2_ concentration-dependent manner (28). The last one is cyAbrB2 binding in the middle or downstream of operons. The middle of *hox*, *pntAB*, and *nifJ* operons and the downstream of *nrt* operon are the cases (Figure 4C). Our data show that genes in the same operon separated by the cyAbrB2 binding region behave differently. In particular, *pntB* is classified as the transiently upregulated gene, while *pntA* is not, despite being in the same operon. This might be explained by the recent study which reported that cyAbrB2 affects the stability of mRNA transcribed from its binding gene (41). The cyAbrB2-mediated stability of mRNA may also account for the decrease in transcript from transient upregulated genes at 4 hours of cultivation. Hereafter, we will focus on the mechanism of the second pattern, regulation by cyAbrB2 on the promoter.

### Insight into the regulation of *hox* and *nifJ* operon by cyAbrB2

Genome-wide analysis indicates that the cyAbrB2-bound region blocks SigE and SigA (Figures 6A and 6B). This is presumably because sigma factors recognize the promoter as a large complex of RNA polymerase. CyAbrB2 binds to the *hox and nifJ* promoter region and may inhibit access to RNA polymerase complex under aerobic conditions. When cells entered microoxic conditions, the boundaries of the cyAbrB2 binding region and cyAbrB2-free region became obscure (Figure 5), and SigE binding peaks on those promoters became prominent (Figure 6C). Notably, cyAbrB2 ChIP efficiency at the *hox* promoter is higher in the microoxic condition than in the aerobic condition (Figure 5). Hence, while the exclusion by cyAbrB2 occupancy on promoter inhibits containing RNA polymerase in the aerobic condition, it is also plausible that chromosomal conformation change governed by cyAbrB2 provides SigE-containing RNA polymerase with access to the promoter region (Figure 8). Our 3C result demonstrated that cyAbrB2 influences the chromosomal conformation of *hox* and *nifJ* region to some extent (Figure 7).

A recent study demonstrated that manipulating the expression of topoisomerase, which influences chromosomal conformational change through supercoiling, affects transcriptional properties in cyanobacteria (15). Moreover, Song et al. pointed out that DNA looping may inhibit transcription in cyanobacteria (41) because artificial DNA looping by the LacI repressor of *E. coli* can repress cyanobacterial transcription (44). Thus, we infer conformation change detected by the present 3C experiment regulates expression of *hox* operon.

### Generality for chromosomal conformation in cyanobacteria

Our 3C analysis revealed that local chromosomal conformation changes upon entry to the microoxic conditions (Figure 8). As cyAbrB2 occupies about 15% of the entire genome and globally regulates gene expression, cyAbrB2 likely governs the whole chromosomal conformation. Furthermore, the conformational changes by deletion of cyAbrB2 were limited, suggesting there are potential NAPs in cyanobacteria yet to be characterized. It is speculated that conformational change of the entire chromosome occurs to deal with many environmental stresses.

### The sigE-mediated mechanism for the transient expression

One possible SigE-mediated mechanism for transient expression is the post-transcriptional activation and degradation of SigE in the dark; i.e., SigE is sequestered by anti-sigma factor under light conditions and released under dark (45), enabling acute transcription of *hox* operon and *nifJ*. Transcripts of *sigE* were continuously downregulated in our time-course transcriptome, while *sigB* (sll0306) and *sigC* (sll0184) were classified as continuous upregulated genes (Supplemental Table S1). It is possible that upregulated SigB and SigC outcompete SigE in prolonged incubation under microoxic conditions. Finally, SigE is degraded under dark within 24 hours (46).

Another reason for the microoxic specific expression may exist in the sequence of the *hox* promoter. We previously determined the consensus sequence of −10 element for SigE regulon in the aerobic condition as “TANNNT,” where N is rich in cytosine (25). The −10 sequence of the *hox* promoter “TAACA<u>A</u>” (23) deviates from the consensus, and no hexamer precisely fitting the consensus is found in the *nifJ* promoter. This deviation can inhibit SigE from binding during aerobic conditions, aside from cyAbrB2-mediated inhibition. Under the microoxic condition, transcription factors LexA (23) and Rre34 (16) may aid SigE binding to the promoter of *hox* and *nifJ*, respectively.

Moreover, SigE seems susceptible to the blocking from cyAbrB2 during the aerobic condition compared with SigA. This is supported by the odds ratio of SigE being in the cyAbrB2-free region was higher than that of SigA in the aerobic condition (Figures 6A and 6B). The higher exclusion pressure of cyAbrB2 on SigE may contribute to sharpening the transcriptional response of hox and nifJ on entry to microoxic conditions. Overall, multiple environmental signals are integrated into the *hox* and *nifJ* promoter through the cyAbrB2 and SigE dynamics.

## Materials and Methods

### Bacterial strains and plasmids

The glucose-tolerant strain of *Synechocystis* sp. PCC 6803 (47) was used as a wildtype strain in this study. The *sigE* (sll1689)-disrupted strain (G50), SigE FLAG-tagged strain, and SigA FLAG-tagged strain were constructed in a previous study (17, 25). Disruption and epitope-tagging of *cyabrb1*(sll0359) and *cyabrb2*(sll0822) were performed by homologous double recombination between the genome and PCR fragment (47). The resulting transformants were selected using three passages on BG-11 plates containing 5 µg/mL kanamycin. Genomic PCR was used to confirm the insertion of epitope tag fragments and gene disruption (Figure S9). Supplemental Tables S2, S3, and S4 contain the cyanobacterial strains, oligonucleotides, and plasmids used in this study.

### Aerobic and microoxic culture conditions

For aerobic conditions, cells were harvested after 24 hours cultivation in HEPES-buffered BG-11_0_ medium (48), which was buffered with 20 mM HEPES-KOH (pH 7.8) containing 5 mM NH_4_Cl under continuous exposure to white light (40 µmol m^−2^s^−1^) and bubbled with air containing 1% CO_2_ (final OD_730_ = 1.4–1.8). For the dark microoxic culture, the aerobic culture cell was concentrated to an OD_730_ of 20 with the centrifuge and resuspended in the culture medium. The concentrated cultures were poured into vials, bubbled with N_2_ gas, and sealed. The sealed vials were shaded and shaken at 30°C for the described times.

### Antibodies and immunoblotting

Sample preparation for immunoblotting was performed as previously described (25), and FLAG-tagged proteins were detected by alkaline-phosphatase-conjugated anti-FLAG IgG (A9469, Sigma Aldrich, St. Louis, MO) and 1-Step NBT/BCIP substrate solution (Thermo Fisher Scientific, Waltham, MA).

### RNA isolation

Total RNA was isolated with ISOGEN (Nippon gene, Tokyo, Japan) following the manufacturer’s instructions and stored at −80°C until use. The extracted RNA was treated with TURBO DNase (Thermo Fisher Scientific) for 1 hour at 37°C to remove any genomic DNA contamination. We confirmed that the A260/A280 of the extracted RNA was >1.9 by NanoDrop Lite (Thermo Fisher Scientific). We prepared triplicates for each timepoint for the RNA-seq library. RT-qPCR was performed as described elsewhere (46).

### ChIP assay

Two biological replicates were used for each ChIP-seq experiment, and one untagged control ChIP was performed. ChIP and qPCR analyses were performed using the modified version of a previous method (25). FLAG-tagged proteins were immunoprecipitated with FLAG-M2 antibody (F1804 Sigma-Aldrich) conjugated to protein G dynabeads (Thermo Fisher Scientific).

### Library preparation and next-generation sequencing

For the ChIP-seq library, input and immunoprecipitated DNA were prepared into multiplexed libraries using NEBNext Ultra II DNA Library Prep Kit for Illumina (New England Biolabs, Ipswich, MA). For the RNA-seq library, isolated RNA samples were deprived of ribosomal RNA with Illumina Ribo-Zero Plus rRNA Depletion Kit (Illumina, San Diego, CA) and processed into a cDNA library for Illumina with the NEBNext Ultra II Directional RNA Library Prep Kit for Illumina (New England Biolabs). Dual-index primers were conjugated with NEBNext Multiplex Oligos for Illumina (Set1, New England Biolabs). We pooled all libraries, and the multiplexed libraries were dispatched to Macrogen Japan Inc. and subjected to paired-end sequencing with HiSeqX. Adapter trimming and quality filtering of raw sequence reads were conducted with fastp (ver. 0.21.0) (49) under default conditions. The paired-end sequences were mapped onto the *Synechocystis* genome (ASM972v1) using Bowtie2 (50) (ver. 2.4.5 paired-end). Supplementary Table S5 contains the read counts that passed via fastp quality control and were mapped by Bowtie2.

### RNA-seq analysis

Mapped reads were counted by HT-seq count (ver. 2.0.2) (51) for the GFF file of ASM972v1, with the reverse-strandedness option. EdgeR package (ver. 3.40.1) (52) was used to perform the differential expression analysis. Fold changes in expression and FDR were used for gene classification. Supplemental Table S6-S8 contains fold change in gene expression calculated by edgeR.

### Genome-wide analyses

Peaks were called using the MACS3 program (ver. 3.0.0b1) (53). For paired-end reads for SigE, SigA, and untagged control ChIP, narrow peaks were called with <1e−20 of the q-value cut-off and “--call-summits” options. The peak summits from two replicates and the untagged control were merged if summits were located within 40 bp of each other. Peak summits identified in both replicates but not in the control, were considered for further analysis. The midpoint of the peak summits for the two merged replicates was further analyzed.

Broad peak calling methods were applied to paired-end reads for cyAbrB2, cyAbrB1, and untagged control ChIP using the “–broad” option, with a q-value cut-off of < 0.05 and a q-value broad cut-off of < 0.05. The intersection of broad peaks from two replicates, excluding those called by the control, was used in subsequent analyses.

The positions of the TSS, including internal start sites, were obtained as reported by (31). The read count, merging, and intersection of the binding region were calculated using BEDTools (ver. 2.30.0) (54). Supplemental Tables S9–S15 contain SigA and SigE peaks and the broad binding regions of cyAbrB2 and cyAbrB1, respectively.

Binding signals in every 100 bp bin for scatter plots were calculated as (IP read counts within 100 bp window) / (input read counts within 100 bp window) * (total input read counts/total IP read counts). GC contents were calculated within 500 bp in 100 bp sliding windows by seqkit (ver. 2.3.0) (55).

### Genome extraction, digestion, and ligation for 3C assay

A 3C assay was conducted based on the previous prokaryotic Hi-C experiment (56, 57), with certain steps modified. To begin, *Synechocystis* were fixed with 2.5% formaldehyde for 15 minutes at room temperature. Fixation was terminated by adding a final concentration of 0.5M of glycine, and cells were stored at −80°C until use. Fixed cells were disrupted using glass beads and shake master NEO (Bio Medical Science, Tokyo, Japan), following the previous study’s instructions for preparing cell lysate for ChIP. The lysates were incubated with buffer containing 1mM Tris-HCl (pH7.5), 0.1mM EDTA, and 0.5% SDS for 10min at room temperature, and 1% Triton X-100 quenched SDS. Genomes in the cell lysate were digested by 600U/mL of HindIII (Takara Bio, Shiga, Japan) for 4 hours at 37°C, and RNA in the lysate was simultaneously removed by 50 µg/mL of RNaseA (Nippon genetics, Tokyo, Japan). The digestion was terminated by adding 1% SDS and 22 mM EDTA. The fill-in reaction and biotin labeling steps were omitted from the procedure. The digested genomes were diluted by ligation buffer containing 1% triton-X100 to the final concentration of approximately 1µg/mL and incubated for 10 min at room temperature. Ligation was performed with 2 U/mL of T4 DNA ligase (Nippon Gene) overnight at 16°C. Crosslinking was reversed under 65°C for 4 hours in the presence of 2.5 mg/mL of proteinase K (Kanto Chemical, Tokyo, Japan), and DNA was purified with the phenol-chloroform method and ethanol precipitation method.

### Preparation of calibration samples for 3C qPCR

Based on a previous study, calibration samples for possible ligated pairs were prepared in parallel with 3C ligation (58). In brief, the purified genome of *Synechocystis* was digested by HindIII, and DNA was purified with the phenol-chloroform and ethanol precipitation. Purified DNA was dissolved into the ligation buffer at a concentration of about 600ng/µL and ligated with 2 U/mL of T4 DNA ligase at 16°C overnight.

### Quantification of crosslinking frequency for 3C assay

Before the real-time PCR assay, we confirmed that each primer set amplified single bands in a ligation-dependent manner by GoTaq Hot Start Green Master Mix (Promega, Madison, WI) (Figure S10). Real-time PCR was performed with StepOnePlus (Applied Biosystems, Foster City, CA) and Fast SYBR Green Master Mix (Thermo Fisher Scientific) according to the manufacturer’s instructions. Interaction frequency was calculated by ΔΔCt method using dilution series of calibration samples described above. We confirmed each primer set amplified DNA fragment with a unique Tm value. The amount of the bait fragment containing *hox* operon were quantified and used as an internal control. Supplemental Table S4 contains the list of primers used in the 3C quantification. Interaction frequency for each primer position was calculated as the relative abundance of ligated fragments against the calibration samples and normalized among samples by internal control.

### Statistical analysis

Statistical analyses were performed with R version 4.2.2 (59). The “fisher.test” function was used for Fisher’s exact test, and p-values < 0.05 were denoted as asterisks in the figure. Multiple comparisons of Fisher’s exact test were conducted using “fisher. Multcomp” function in the RVAideMemoire package (60), where p-values were adjusted by the “fdr” method and FDRs <0.05 are shown in the figures. Multiple comparisons of the Wilcoxon-rank test were conducted by “pairwise.wilcox.test,” and p-values were adjusted by the “fdr” method. Adjusted p-values < 0.05 are shown in the figure. The correlation coefficient was calculated with the “cor” function. Gene set enrichment analysis (GSEA) was performed by culsterPlofiler package (61) in R with p-value cut-off of 0.05. The enriched pathways detected by GSEA are listed in Supplemental Table S16.

### Manuscript writing

We employed GPT-3.5 to refine the expression of the manuscripts.

### Accession Numbers

Raw ChIP sequencing and RNA sequencing reads were deposited in the Sequence Read Archive (accession ID: PRJNA956842).

## Supporting information

Supplemental Talbles5-16

## Acknowledgments

This study was supported by the following grants to T.O.: Grant-in-Aid for Scientific Research (B) (grant no. 20H02905), JST-ALCA of the Japan Science and Technology Agency (grant number JPMJAL1306), the Asahi Glass Foundation, and JST-GteX (grant number JPMJGX23B0). We thank Dr. Kohki Yoshimoto for providing laboratory instruments and Ms. Kaori Iwazumi for the support of bacterial culture and the medium preparation.

## Author contributions

R.K. and T.O. designed the study; R.K. conducted the experiment; R.K. analyzed the data; R.K. and T.O. wrote the paper.

---

**Figure S1.**
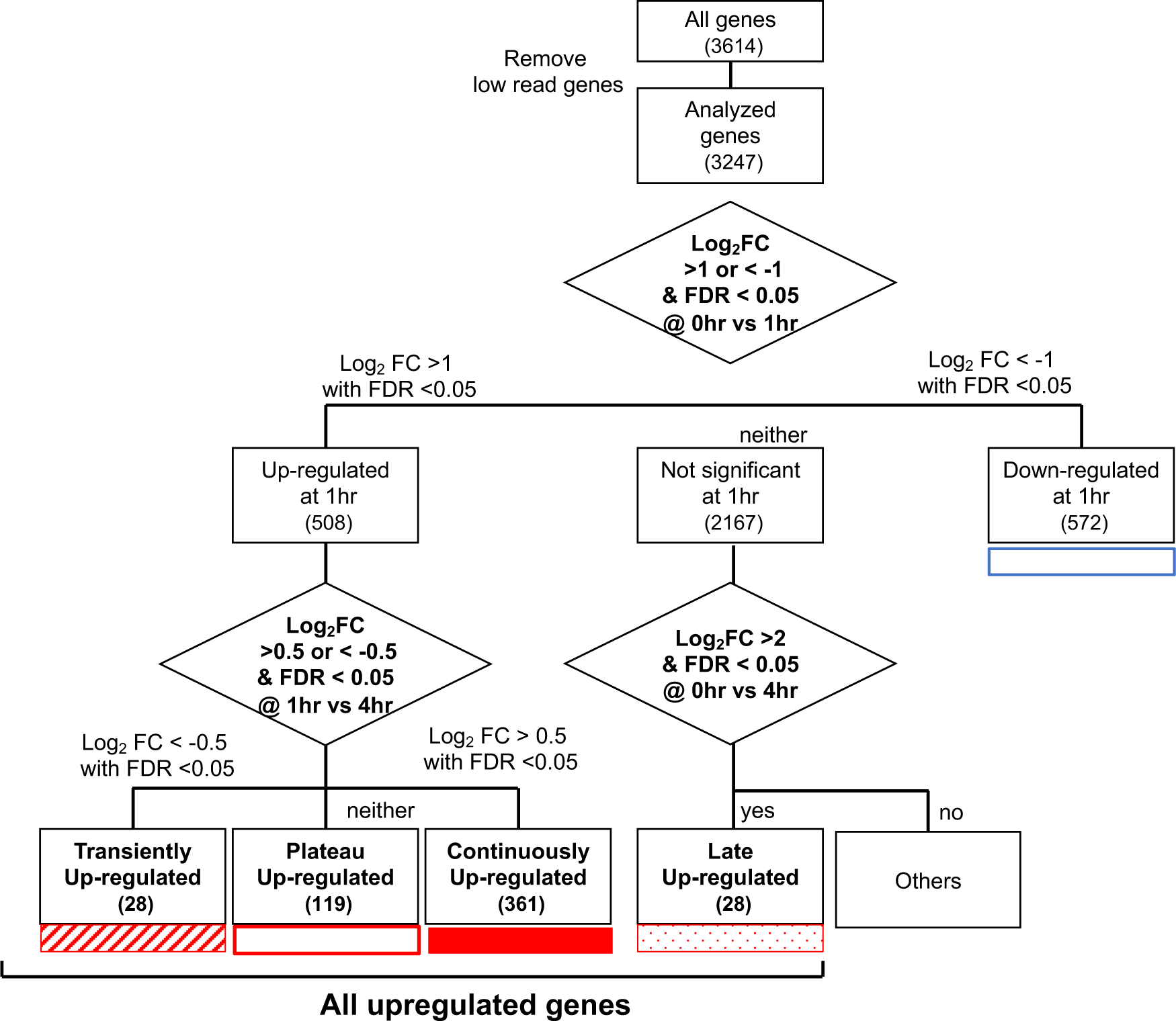
Schematic diagram showing the classification of genes according to the time-course transcriptome. Transient (striped square), Plateau (open square), Continuous (filled square), and Late (dotty square) are denoted as all upregulated genes.

**Figure S2.**
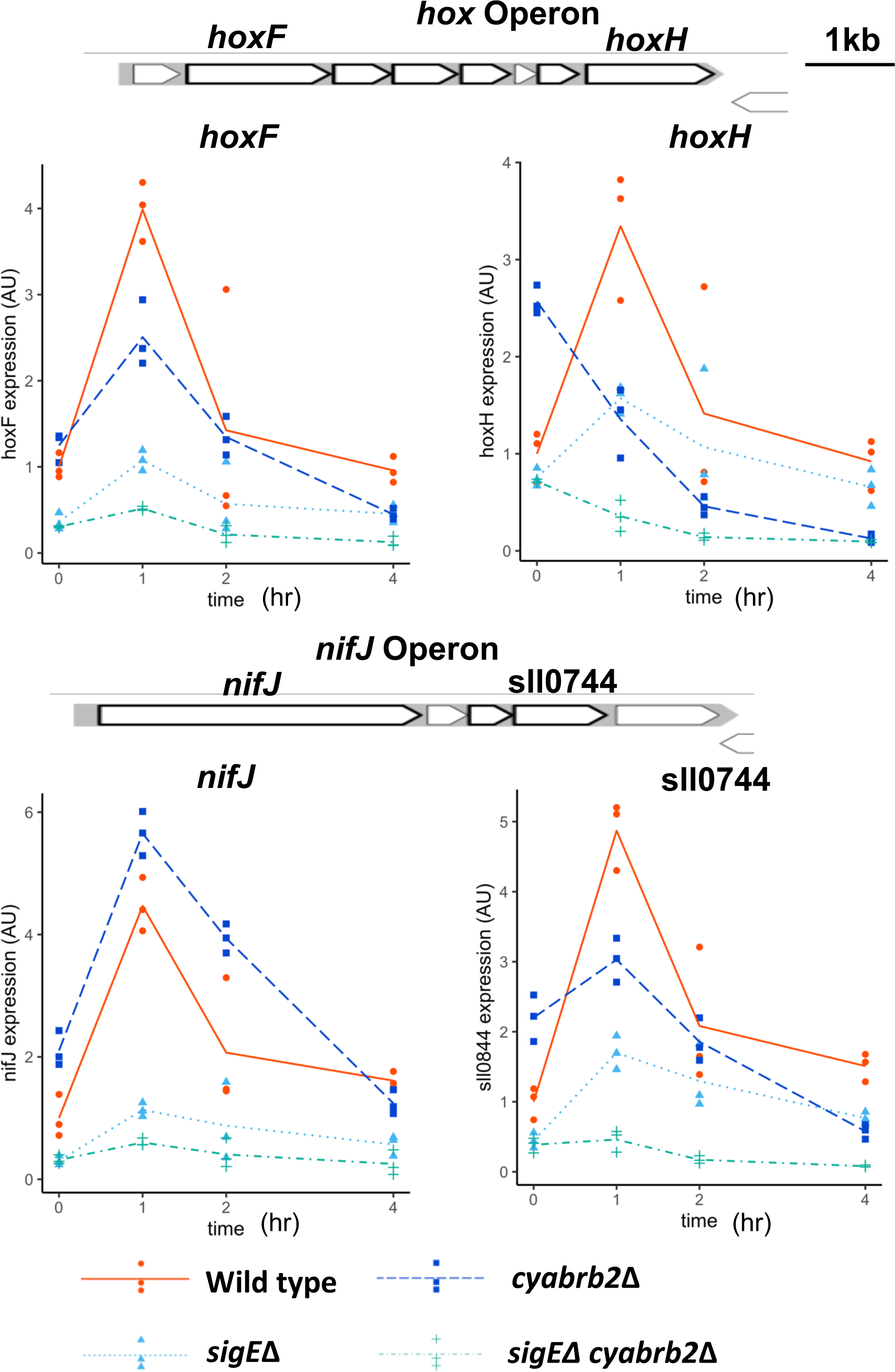
RT-qPCR validated the transiently upregulated genes classified by RNAseq. Transcripts extracted from wildtype (solid line), Δ*sigE* mutant (dotty line), Δ*cyabrb2* mutant (dashed line), and Δ*sigE* Δ*cyabrb2* double mutant (dot-dashed line) were assayed in the aerobic condition (0hr) and 1,2,4 hour incubation of microoxic conditions. As the representative of the transiently upregulated genes, expression of *hoxF*, *hoxY, nifJ,* and *sll0744* were quantified by RT-qPCR. The line represents the mean of n=3, and individual data points are shown as dot plots. Data of each gene is normalized by the mean score of wildtypes in the aerobic condition.

**Figure S3.**
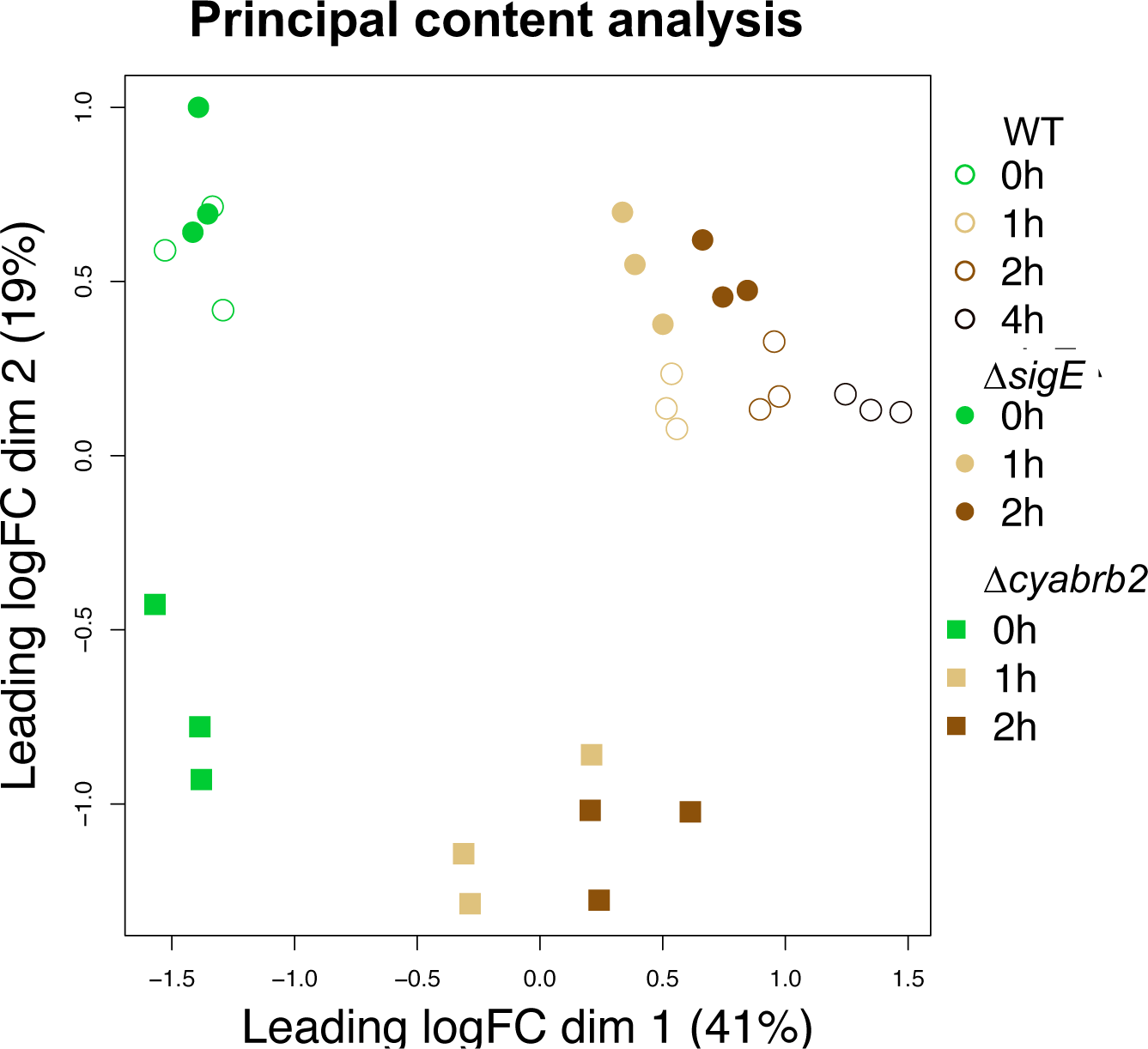
Primary component scatter plot showing the profiles of RNA-seq data.

**Figure S4.**
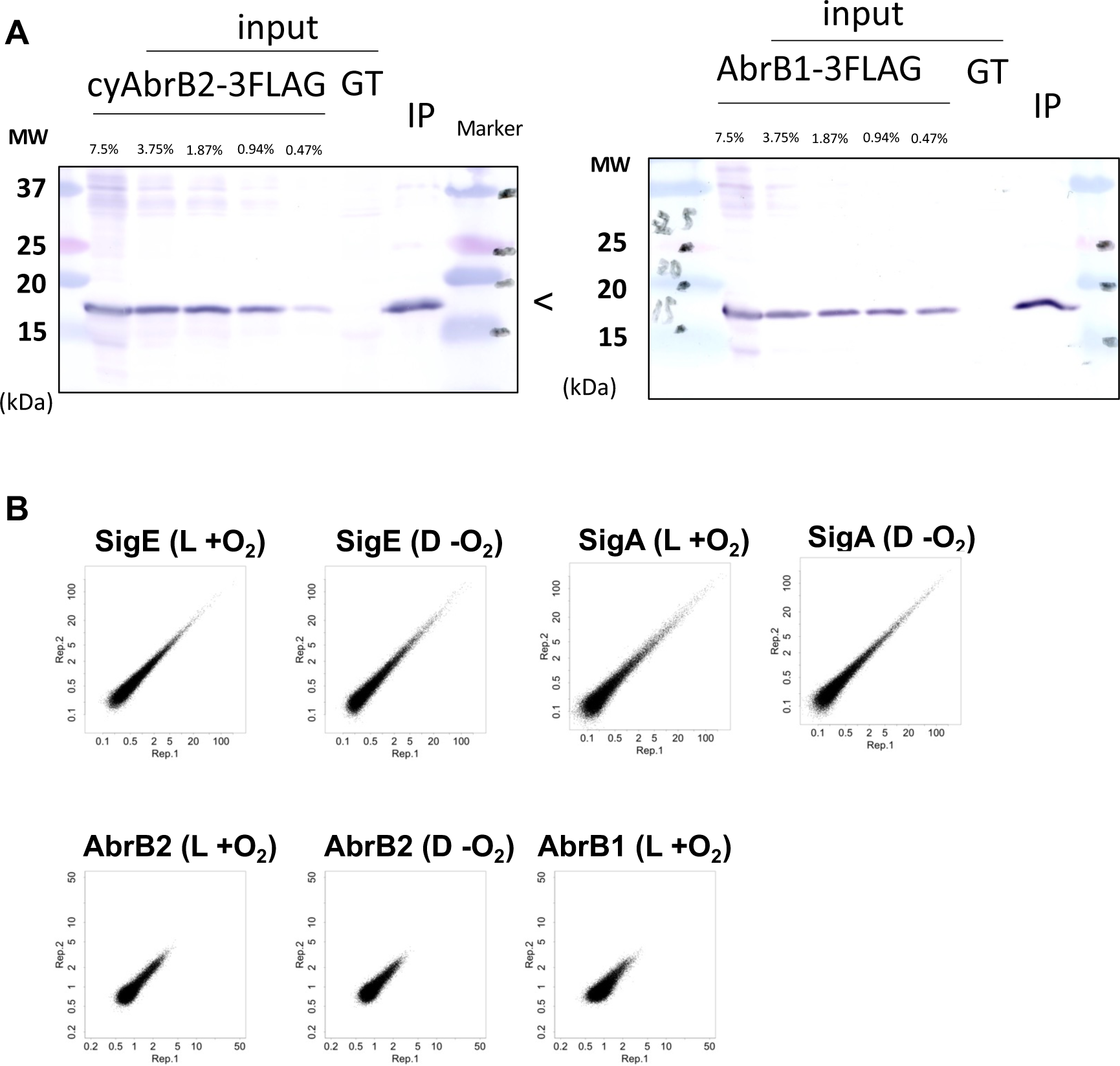
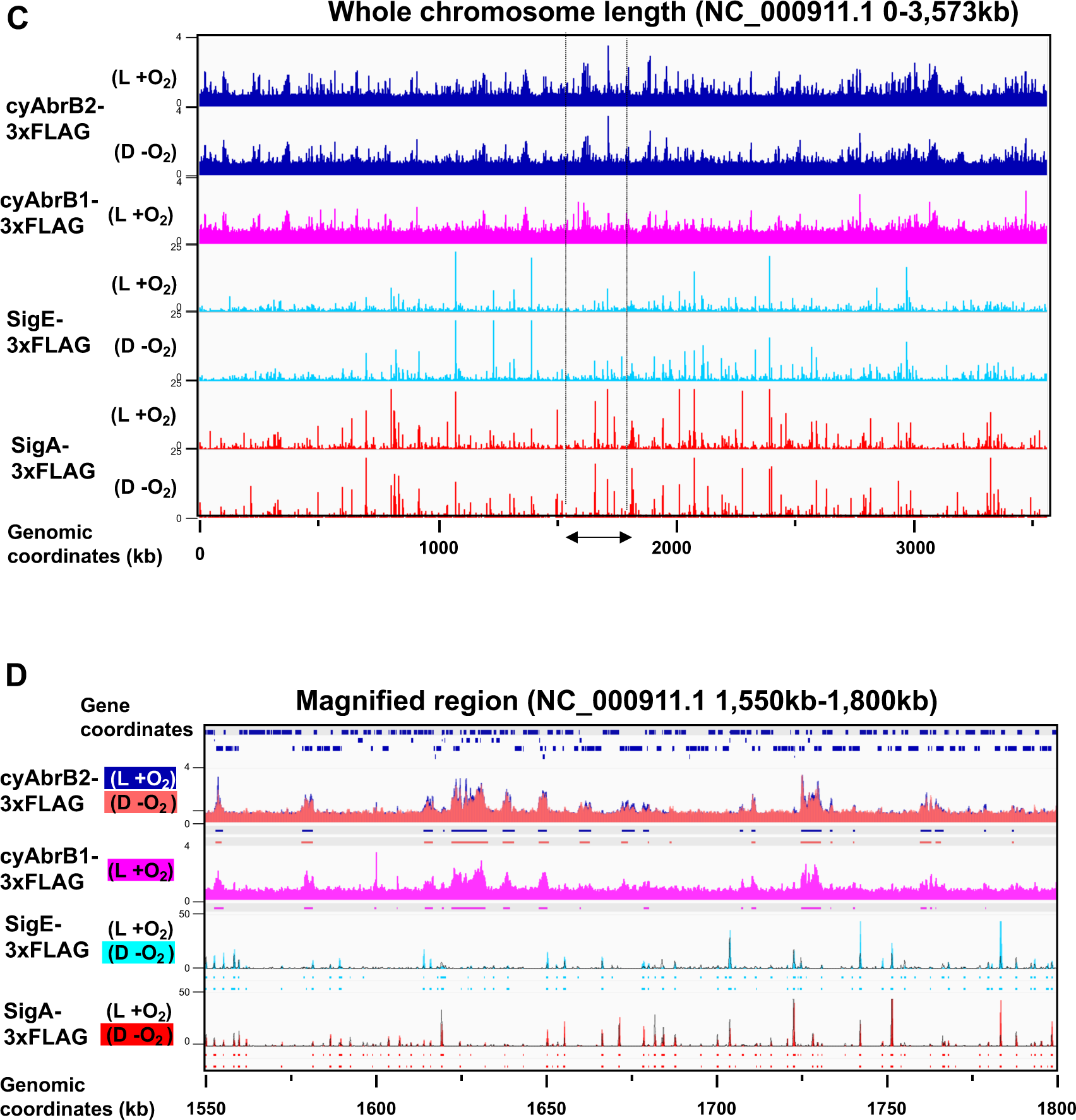
The procedure for ChIP-seq of flag-tagged proteins and Overview of ChIP-seq data. Validation of procedure for ChIP-seq of flag-tagged cyAbrB2, SigE, and SigA. (A) The immunoblot for inputs and immunoprecipitants (IP) of ChIP for cyAbrB2-FLAG and cyAbrB1-FLAG. Input lysate of untagged control (GT) is also loaded. Inputs equivalent to the indicated portion of IP were loaded. (B)Scatter plots showing the reproducibility of two replicates for ChIP-seq assay. ChIP-seq data of SigE, SigA, and cyAbrB2 in aerobic and microoxic conditions and ChIP-seq data of cyAbrB1 in the aerobic condition are shown. Dots indicate normalized IP read count / normalized input read count in each 100bp window. X-axis is the value of replicate1, and Y-axis is the value of replicate 2. (C) and (D) Overview for ChIp-seq of flag tagged cyAbrB2, cyAbrB1, SigE, and SigA. Y-axis indicates [ normalized IP read count / normalized input read count at each 25bp window], and X-axis indicates chromosome position. (C) Distribution of cyAbrB2, cyAbrB1, SigE, and SigA across the whole genome of *Synechocystis*. Aerobic (L +O_2_) and dark microoxic (D -O_2_) data are displayed. (D) Magnified image for chromosome position of 1,550kb-1,800kb. ChIP-seq data of cyAbrB2, cyAbrB1, SigE, and SigA in aerobic and dark microoxic conditions are overlayed. The dots below the graph indicate the binding region of each protein calculated by peak caller.

**Figure S5.**
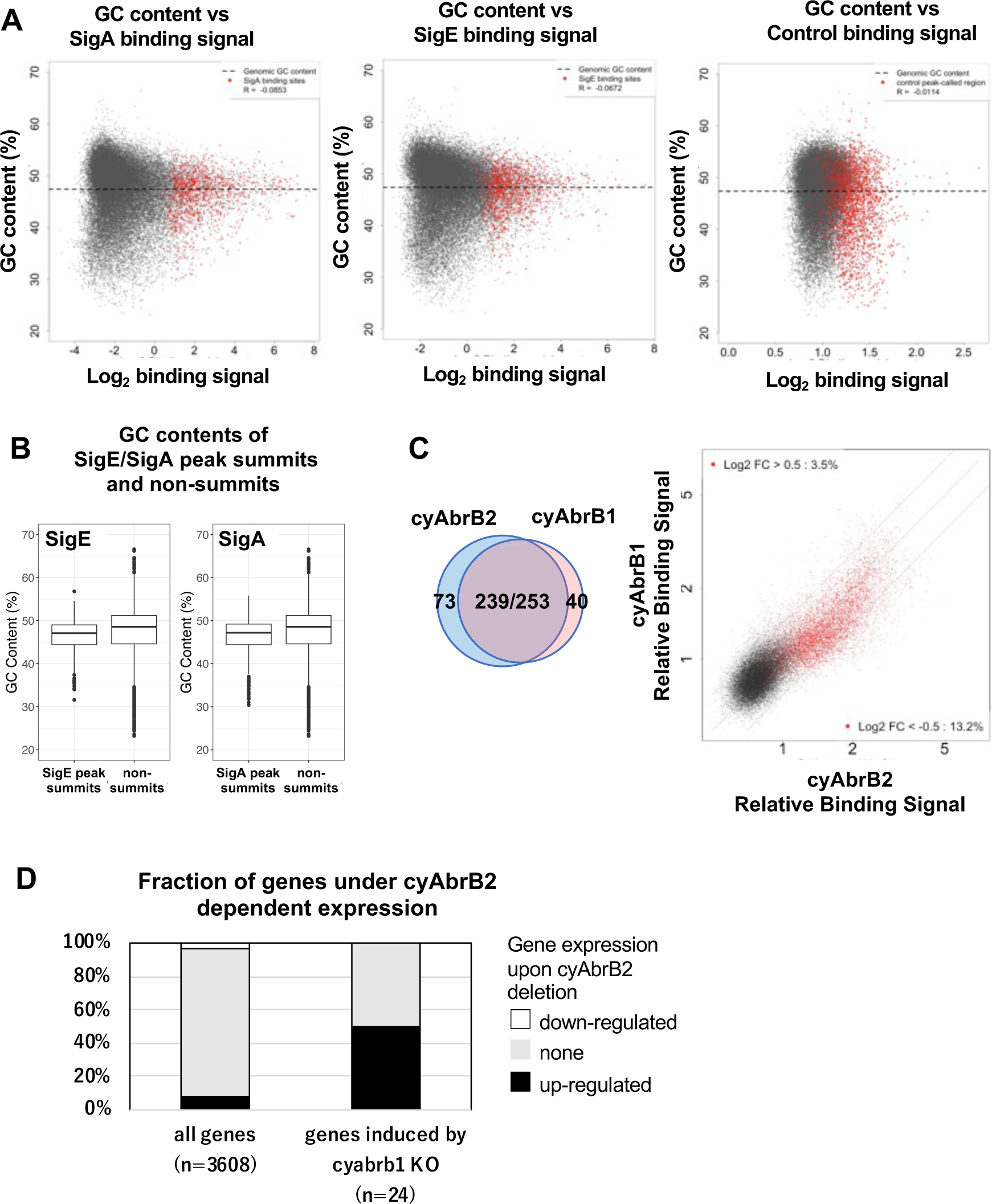
GC content vs ChIP enrichment score of SigA and SigE. (A) Scatter plot showing GC contents in each 100bp vs. binding signal of SigA, SigE, and control IP. Data are displayed as in Figure 3C. (B) GC content in each 100bp of (left) SigE peaks and non-SigE peaks, and (right) SigA peaks and non-SigA peaks. (C) Venn diagram showing overlap of the binding region of cyAbrB1 and cyAbrB2 (left), and scatter plot showing ChIP binding signal of cyAbrB2 (y-axis) and cyAbrB1(x-axis) in the aerobic condition. Data is plotted as in Figure 5A. (D) Fractions of upregulated and downregulated genes upon the Δ*cyabrb2* mutant in the aerobic conditions. Fractions of all genes (left n=3608) and genes induced by cyAbrB1 knockout (right n=24) are shown. Genes induced by cyAbrB1 knockout are from the previous study (30).

**Figure S6.**
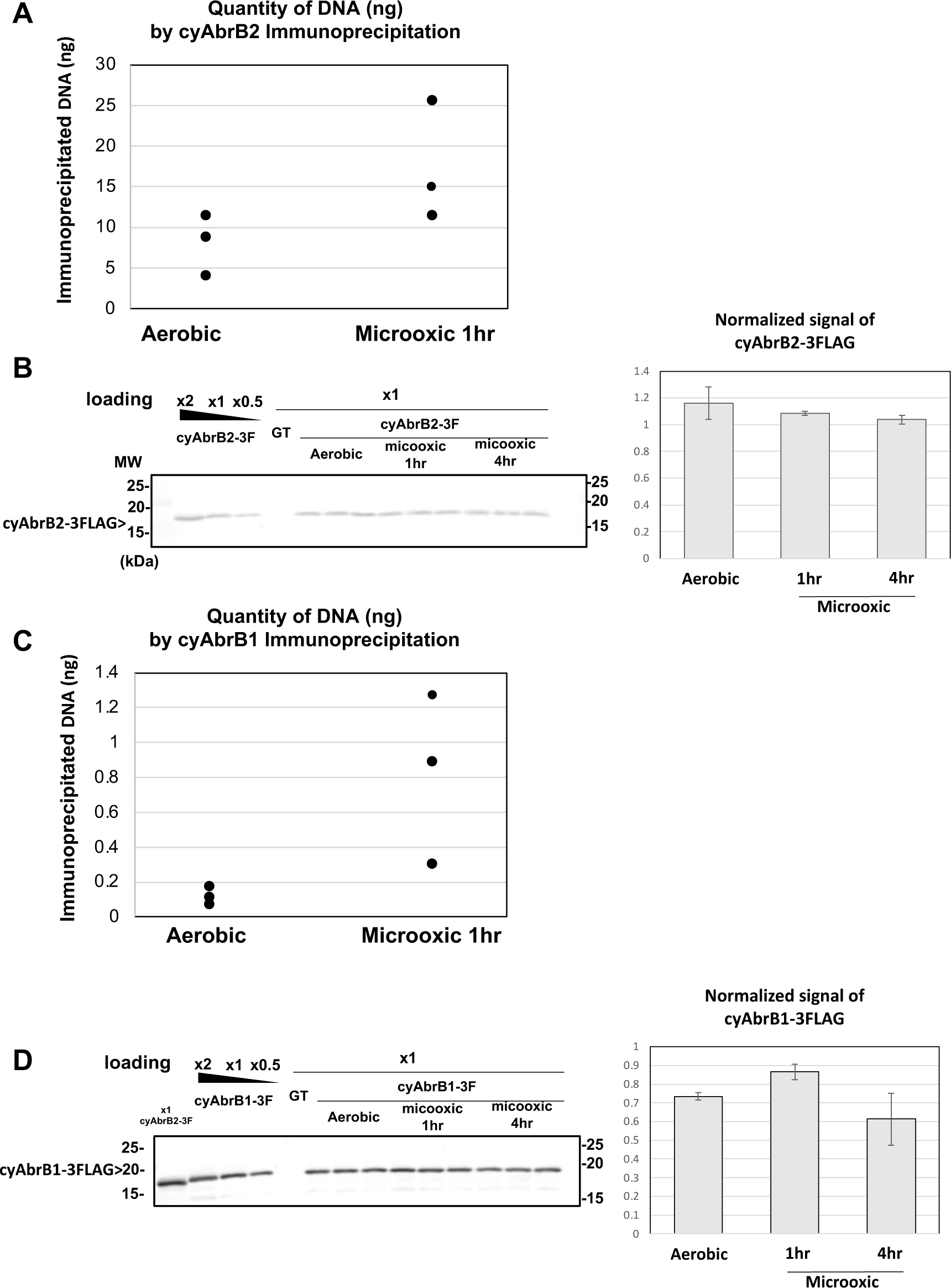
Alteration of cyAbrB2 binding to genome under the microoxic condition. (A) and (C) Amount of precipitated DNA by cyAbrB2 ChIP and cyAbrB1 ChIP, respectively. Three experiments were performed in the aerobic and microoxic conditions. (B) and (D) Western blot images of cyAbrB2-3FLAG and cyAbrB1-3FLAG, respectively. Proteins were extracted in the aerobic condition and 1hr and 4hr incubation under microoxic conditions. The total protein concentration of each sample was adjusted to 4mg/mL, measured by the BCA method. Quantification of cyAbrB2 from western blot image was performed by ImageJ (ver. 2.0.0-rc-65) and plotted in the right graph.

**Figure S7.**
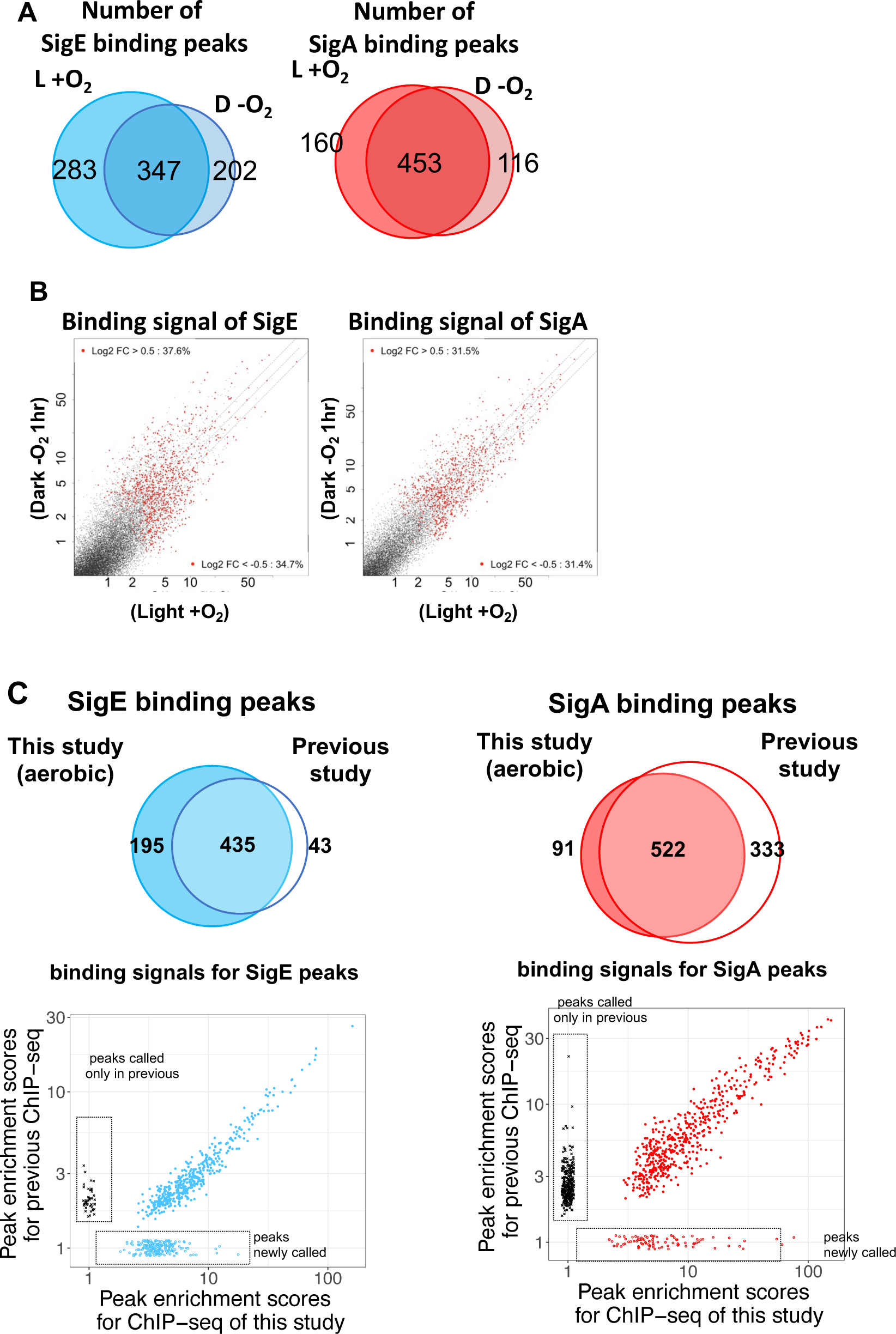
(A) Venn diagram showing the number of peaks of SigE (left) and SigA (right) in aerobic (L + O_2_) and dark microoxic (D − O_2_) conditions. (B) Scatter plot showing changes in the binding signal of SigE and SigA by 1 hour cultivation under microoxic conditions. The binding signal of each 100 bp window is plotted. (C) Reproducibility of ChIP-seq data of SigA and SigE, compared with the previous study (Kariyazono and Osanai 2022). (top) Venn diagrams show the overlapping of peaks called in this study and the previous study. (bottom) Scatter plot comparing ChIP binding signals of SigA and SigE peaks commonly called in present and previous studies. Plots boxed by dashed lines are peaks called only in the present or previous study.

**Figure S8.**
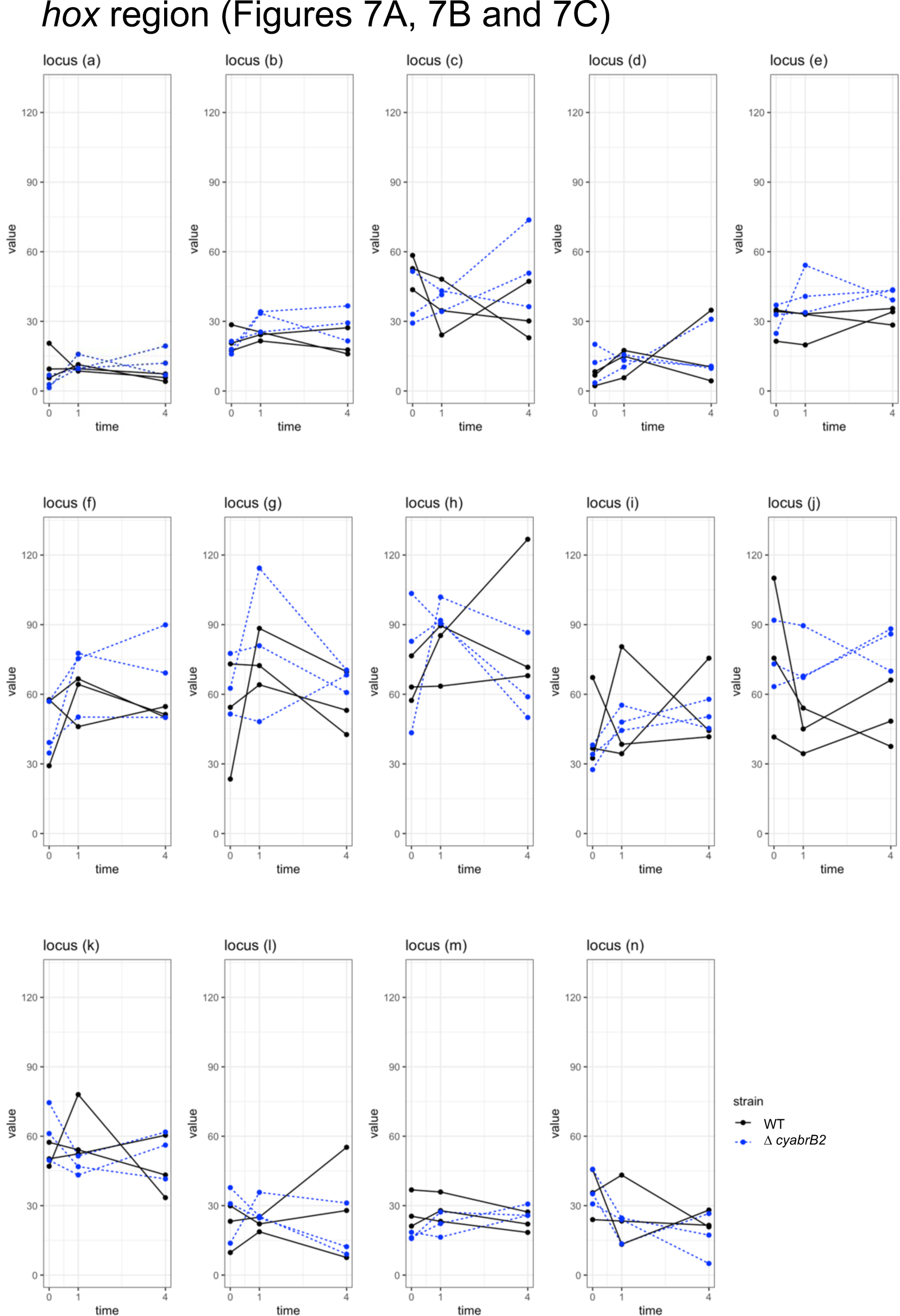
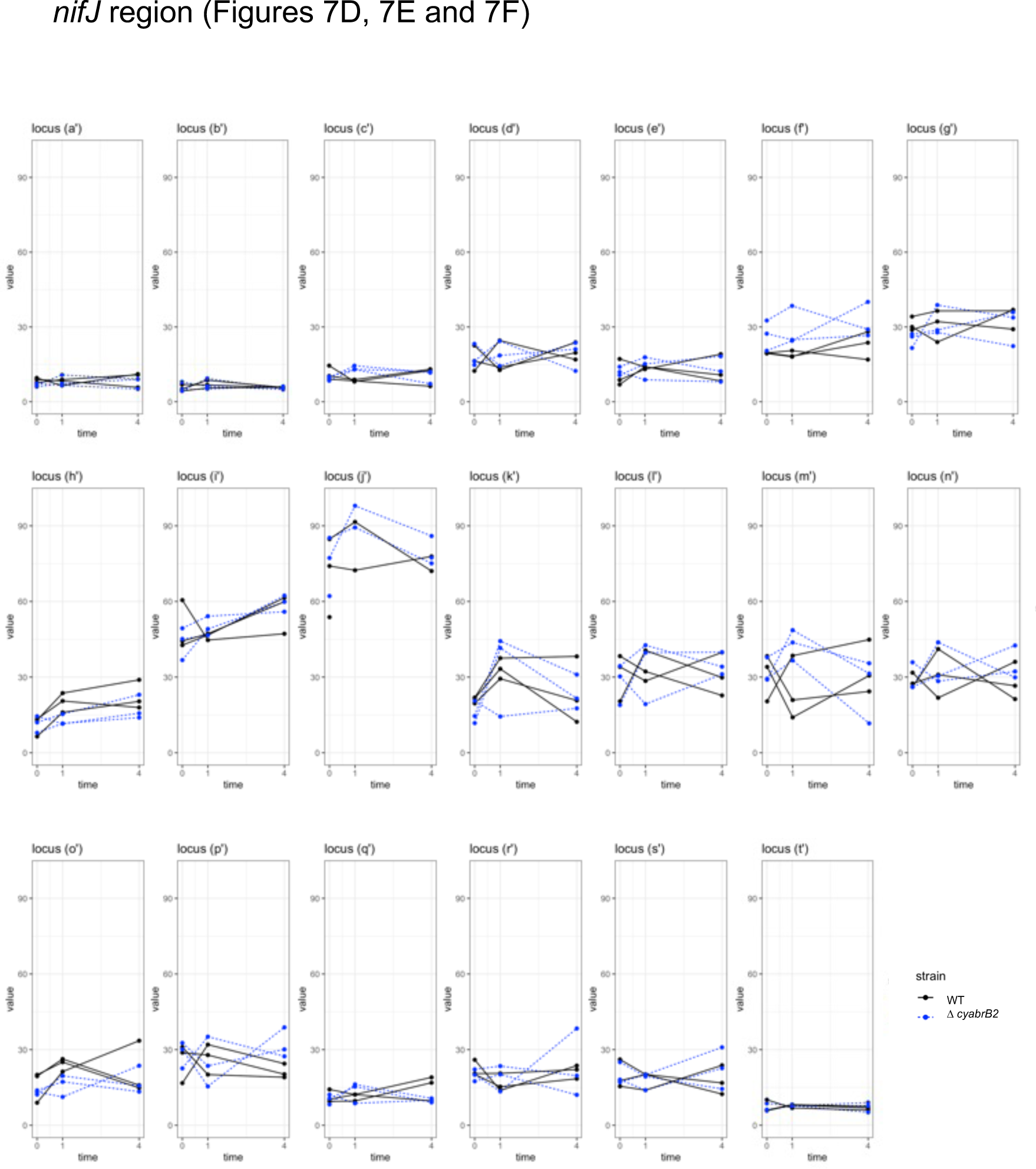
Dynamics of individual 3C scores. Re-plotting of Figure 5B and 5C with the x-axis showing time (0,1,4 hr in microoxic conditions) and the y-axis showing the interaction frequency. Plots from the individual samples are connected by solid (Wildtype) or dotty (Δ*cyabrb2*) lines.

**Figure S9.**
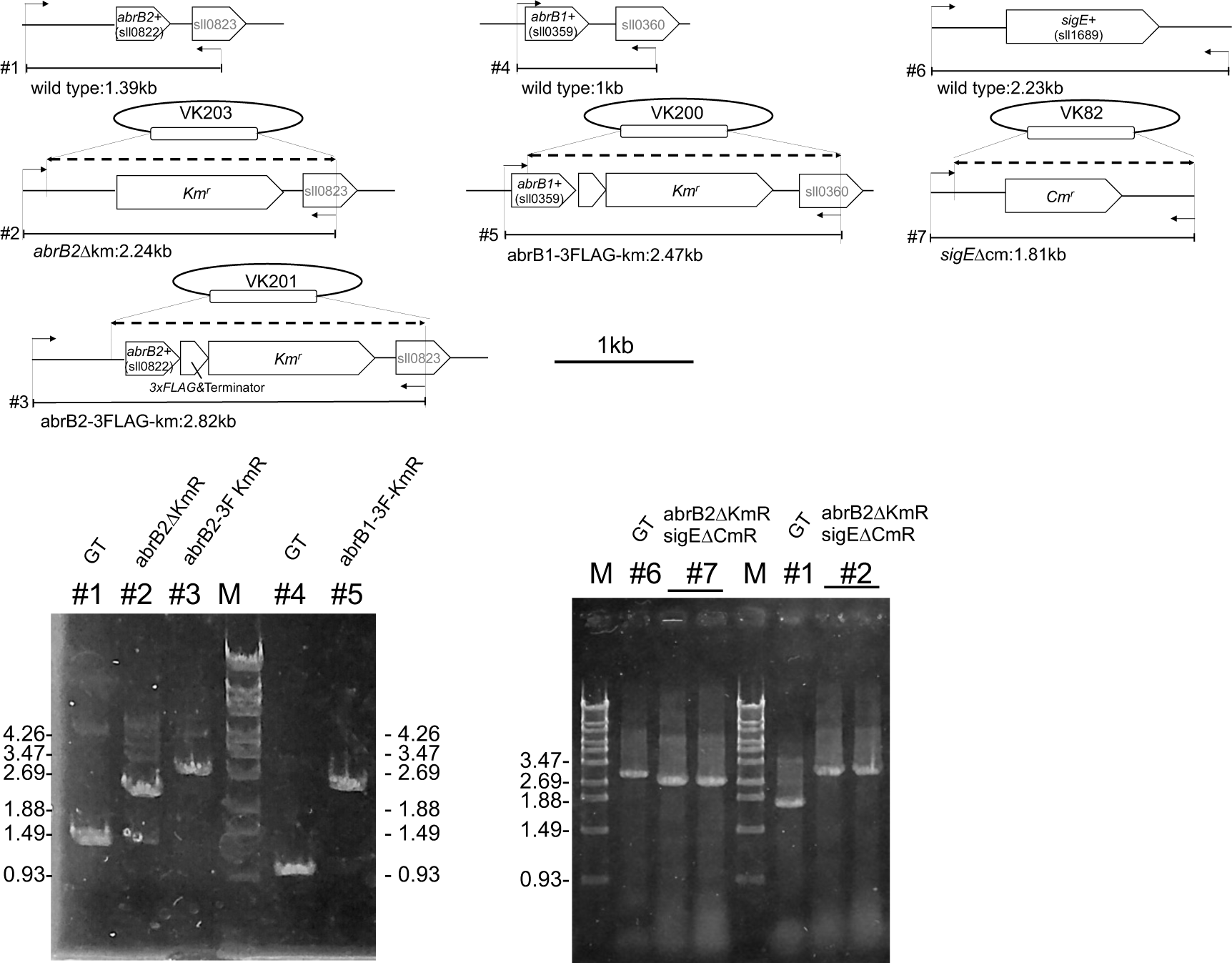
Confirmation of genomic deletion and the epitope tagging of *abrB2* (#1-#3), the epitope tagging of *abrB1* (#4 and #5), and deletion of *sigE* (#6 and #7). Dotty lines are homologous regions between plasmids and the genome. Arrows indicate the position of check primers.

**Figure S10.**
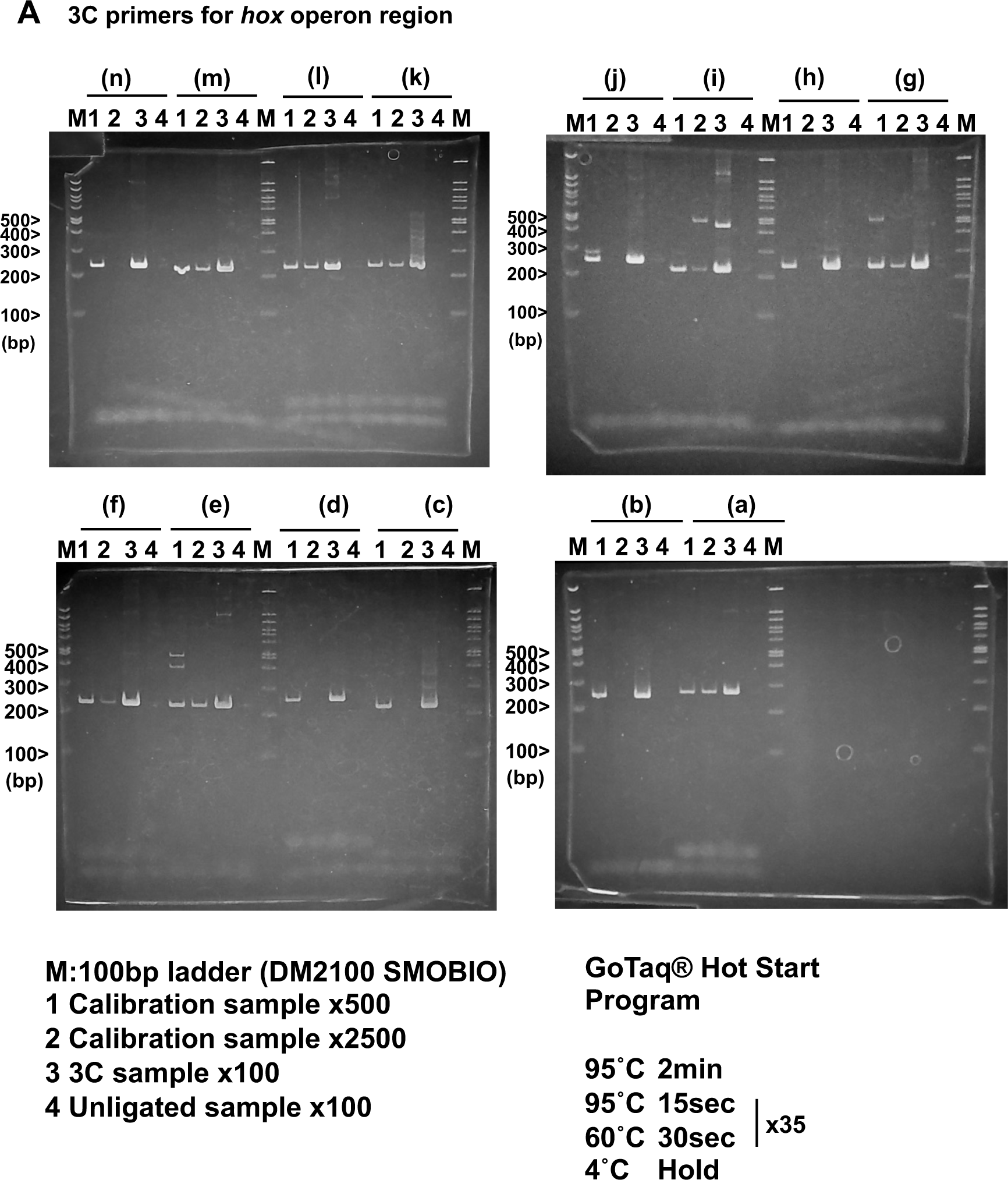
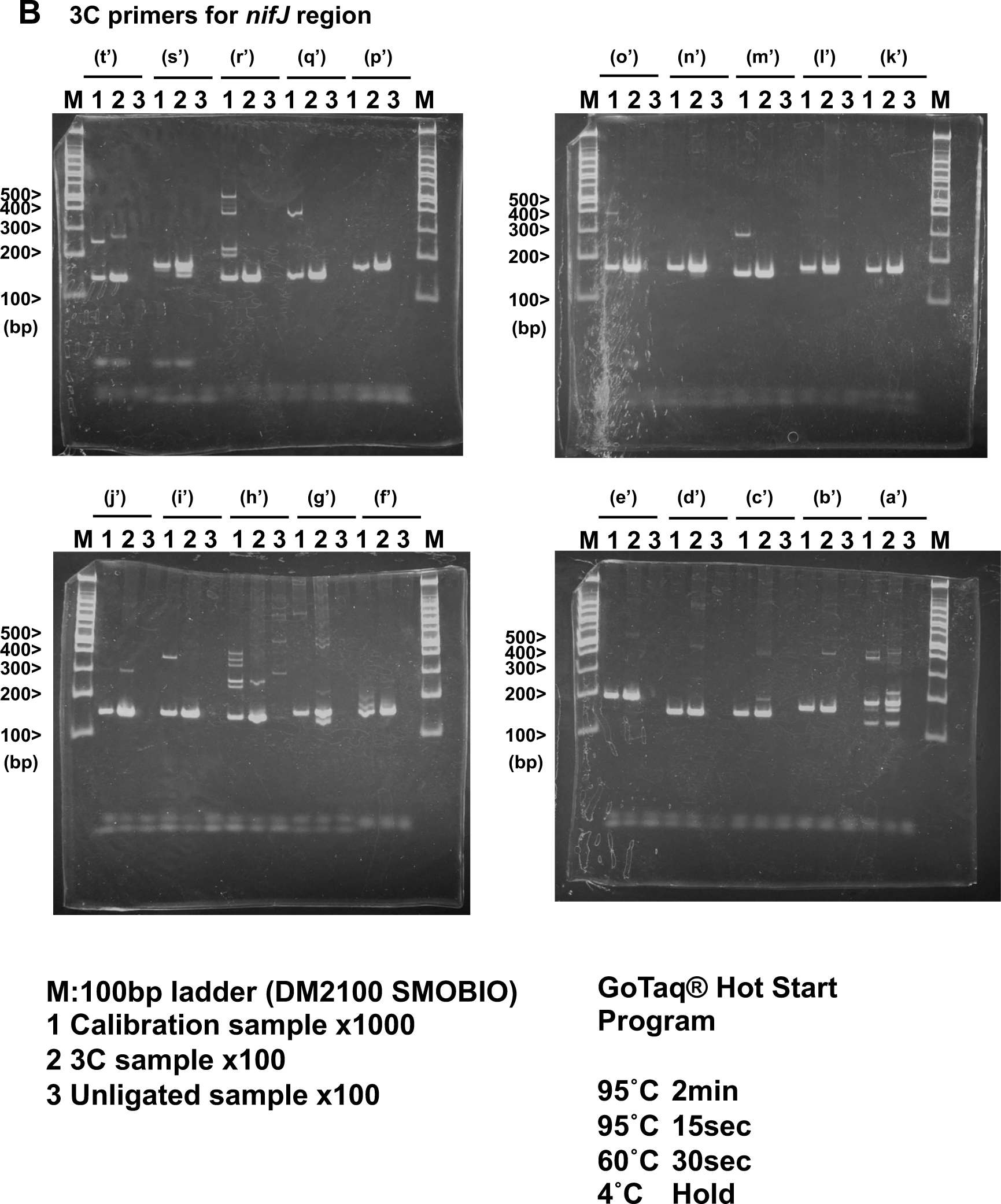
Validation of unidirectional primer sets for 3C assay. The validation of unidirectional primer sets for 3C assay is shown in Figure 7. The 3C sample in this assay is the mixture of all 3C samples assayed in Figure 7.

**Table S1.** Fold changes of transcripts from *sigA, sigB, sigC, sigD, and sigE*.

| Sigma factor | locus | 0hr vs 1hr |  | 1hr vs 4hr |  |
| --- | --- | --- | --- | --- | --- |
|  |  | Log2FC | FDR | Log2FC | FDR |
| SigA | slr0653 | -0.873248 | 0.00766486 | -0.0013514 | 0.99797563 |
| SigB | sll0306 | 1.38098826 | 8.42E-06 | 0.77453605 | 0.04057775 |
| SigC | sll0184 | 2.97101055 | 1.75E-16 | 1.30743549 | 0.00067892 |
| SigD | sll2012 | 0.4701823 | 0.1498473 | -0.4522181 | 0.32402556 |
| SigE | sll1689 | -1.9111759 | 1.96E-11 | -1.1223298 | 0.00633142 |

Data is extracted from Supplemental Table S7.

**Table S2.** List of strains used in this study.

| Strain ID | Host strain | genotype | Reference |
| --- | --- | --- | --- |
| GT |  |  | Williams, 1988 |
| G50 | GT | $\Delta sigE::KmR$ | T. Osanai, et al,2005 |
| KR93 | GT | SigA-8His-KmR | (Kariyazono and Osanai 2022) |
| KR94 | GT | SigA-3FLAG-KmR | (Kariyazono and Osanai 2022) |
| KR340 | GT | $\Delta cyabrB2::KmR$ | in this study |
| KR338 | GT | cyAbrB(sll0359)-3xFLAG-KmR | in this study |
| KR339 | GT | cyAbrB2(sll0822)-3xFLAG-KmR | in this study |
| KR359 | KR340 | $\Delta cyabrB2::KmR \Delta sigE::CmR$ | in this study |

**Table S3.**
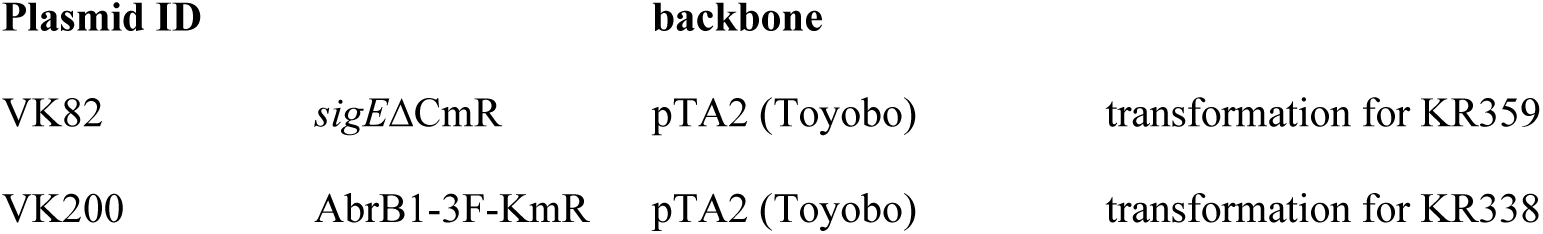
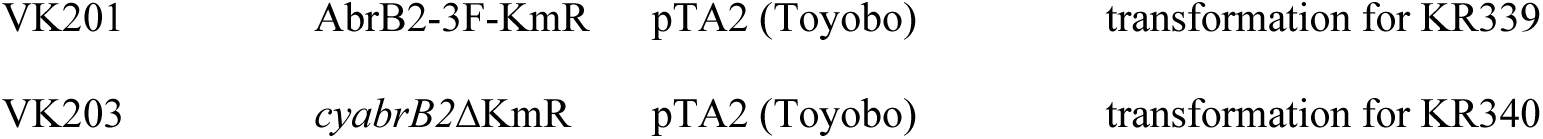
plasmids used in this study.

**Table S4.**
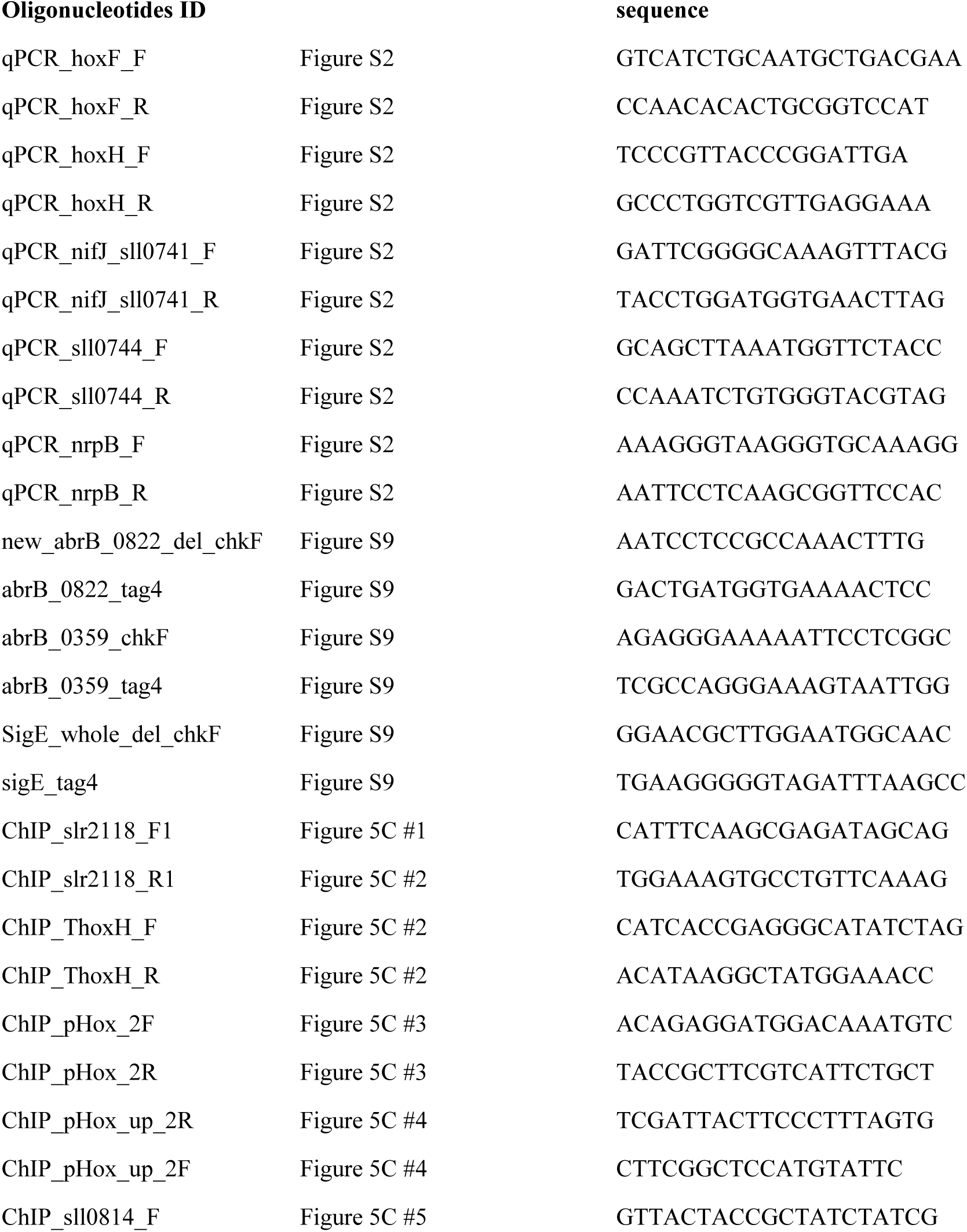
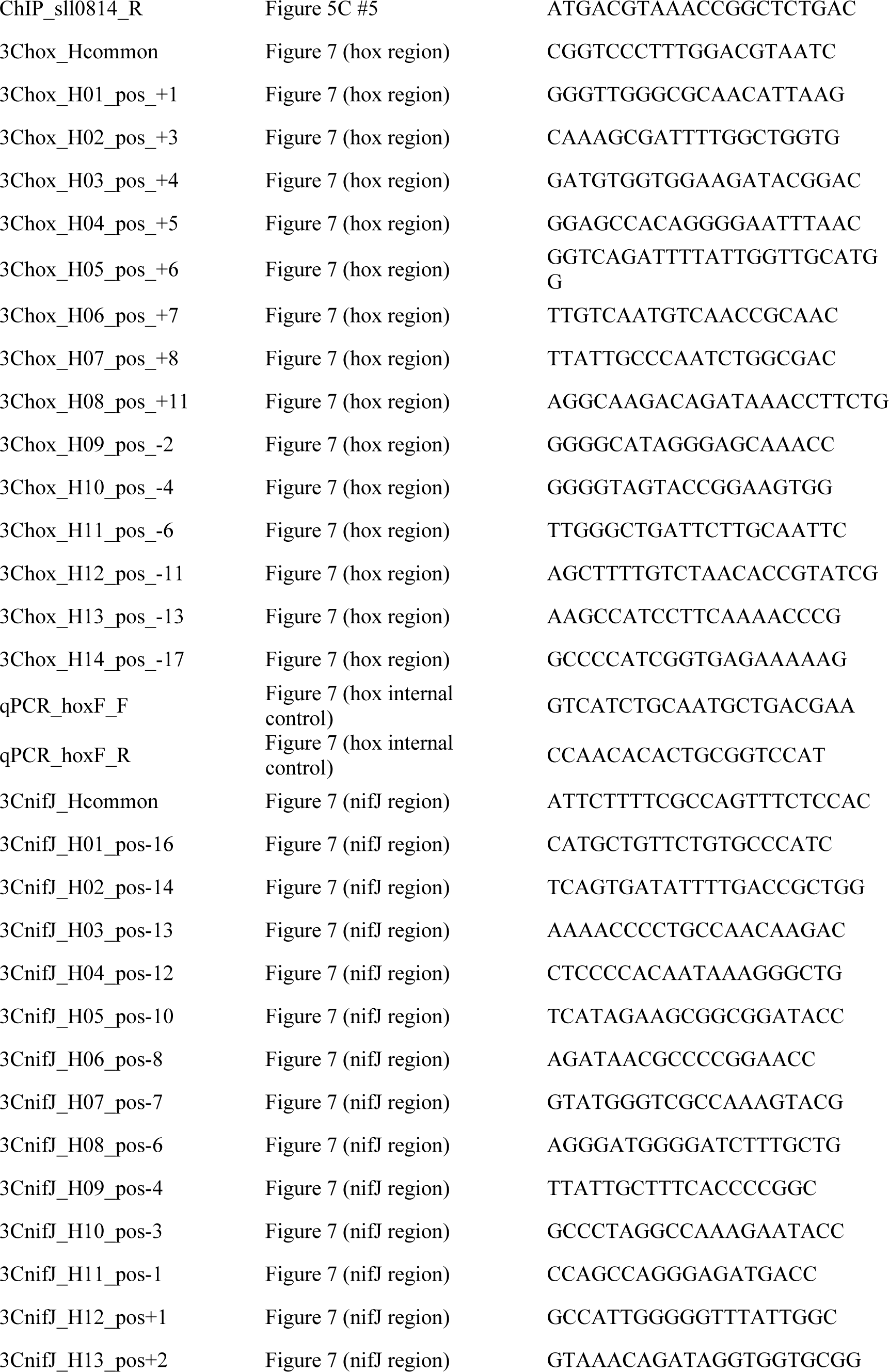
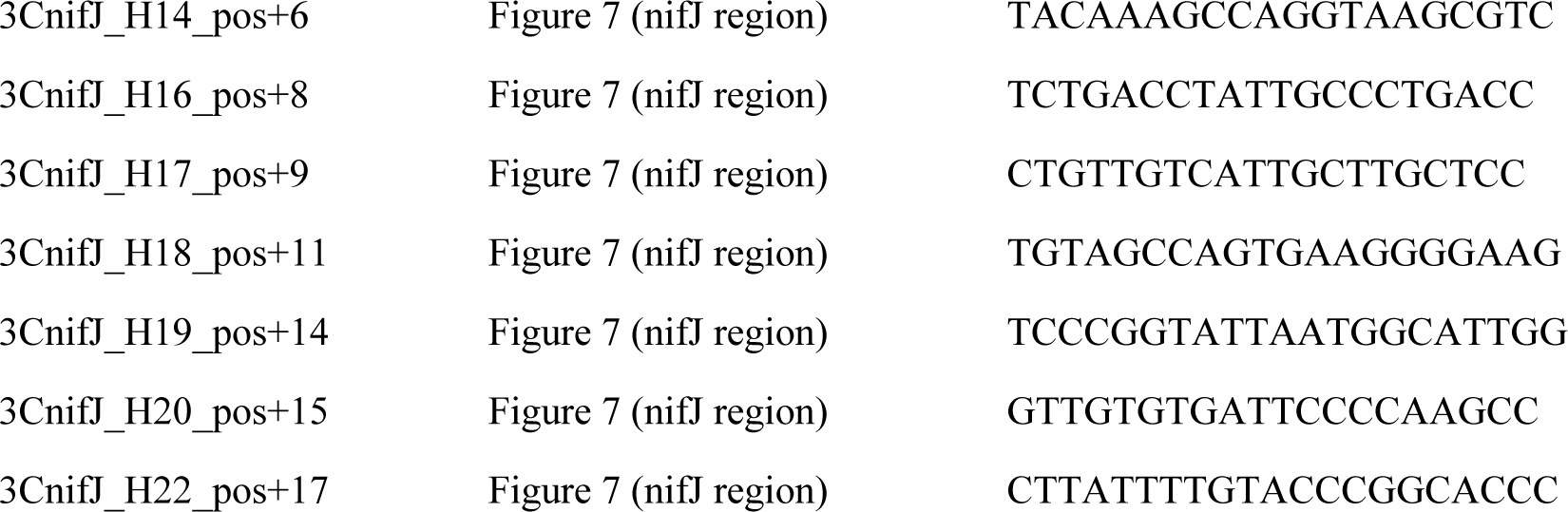
oligonucleotides used in this study.

